# A single cortical circuit implements fast and slow visual search

**DOI:** 10.64898/2026.09.09.750509

**Authors:** Mattia F. Pagnotta, Haojun Zhuang, S. J. Katarina Slama, David King-Stephens, Kenneth D. Laxer, Tor Endestad, Anne-Kristin Solbakk, Jack J. Lin, Robert T. Knight

**Affiliations:** Helen Wills Neuroscience Institute, University of California Berkeley, USA; Department of Neurology, University of California Irvine, USA; California Pacific Medical Center, San Francisco, USA; University of Oslo, Norway; Helgeland Hospital, Mosjøen, Norway; Oslo University Hospital – Rikshospitalet, Norway; Department of Neurology, University of California Davis, USA; Departments of Psychology and Neuroscience, University of California Berkeley, USA

## Abstract

How the brain produces different behaviors from a fixed anatomical substrate is a foundational question in neuroscience. The dominant account holds that behavioral flexibility requires distinct neural circuits: dedicated systems selectively recruited for different processing modes. An alternative proposes that flexibility arises not from switching between circuits, but from dynamically controlling the speed at which a shared circuit operates. Here, we test this account directly using intracranial EEG recordings from nineteen human patients performing two visual search tasks of markedly different difficulty: an easy color-singleton search and a difficult orientation search, a fast vs. slow dissociation attributed for four decades to distinct anatomical and oscillatory neural systems. We show that both search tasks recruit the same cortical regions, oscillatory frequencies, and processing hierarchy. Dynamic time warping reveals that the two neural trajectories are time-stretched versions of one another; the same temporal scaling appears within each condition between faster and slower trials, indexing response speed rather than the distinction between tasks. Alpha-band oscillations (∼8–10 Hz) support this temporal scaling through two coordinated mechanisms: synchronizing activity across a posterior cortical network and phase-gating local high-frequency activity (70–150 Hz) within those regions at the same preferred phase in both conditions. These findings establish that, for the two tasks studied here, fast and slow visual search are not implemented by selecting between distinct dedicated circuits, but by adjusting the speed of a single alpha-coordinated architecture. The same principle may apply to other search tasks, and to other domains in which dual-process architectures have been invoked, a testable prediction for future studies.

## 1. Main

The brain routinely produces different behaviors from a fixed anatomical substrate. A surgeon’s hand executes the same reaching movement with fine deliberate precision or rapid automatic skill depending on context; a reader processes the same sentence slowly when encountering a new argument or effortlessly when skimming a familiar one; a forager detects a familiar object instantly in a cluttered scene or searches it out methodically when it is camouflaged. This behavioral flexibility, the ability to shift between processing modes without rewiring the brain, is one of the defining computational achievements of the human nervous system, and one of its least understood. Two broad classes of neural solutions have been proposed. In the “circuit-segregation” account, distinct specialized systems are selectively recruited depending on task demands: flexible behavior reflects the selection of the appropriate circuit from a repertoire of dedicated modules (Corbetta & Shulman, 2002; Fodor, 1983). In the “circuit-dynamics” account, the same anatomical substrate generates different behavioral modes through reconfiguration of its temporal or oscillatory coordination: flexibility is a property of neural dynamics, not of anatomy (Buzsáki & Draguhn, 2004; Fries, 2005; Varela et al., 2001). These two accounts make fundamentally different predictions about the neural architecture of cognition, yet distinguishing between them has proved difficult. Neuroimaging reveals where activity occurs but cannot resolve the millisecond-scale dynamics that coordinate it (Logothetis, 2008). Electrophysiology resolves dynamics but typically samples too sparsely to map the spatial architecture of active circuits (Cohen, 2014, 2017). What is needed is a method that simultaneously characterizes both the spatial map and the temporal dynamics of active networks, and a behavior that is well characterized, which has developed competing theories that can be directly tested.

Visual search offers precisely this opportunity. For four decades it has provided one of the clearest behavioral dissociations in cognitive neuroscience: targets defined by a single feature (a red item among green ones) appear to “pop out” of the display with reaction times independent of the number of distractors, while targets defined by a conjunction of features (a red vertical bar among red horizontals and green verticals) require serial, attention-demanding search where reaction times increase steeply with set size (A. M. Treisman & Gelade, 1980; A. Treisman & Sato, 1990; Wolfe, 2014, 2021). This ’Pop-out’ versus ’Search’ dissociation has been attributed, historically, to two distinct neural systems: a bilateral dorsal attention network (DAN) supporting goal-directed, top-down attentional deployment in Search, and a right-lateralized ventral network (VAN) supporting stimulus-driven salience detection in Pop-out (Corbetta et al., 2008; Corbetta & Shulman, 2002). Single-unit and local field potential recordings in macaque prefrontal and parietal cortex extended this two-systems account to the oscillatory level, showing that top-down Search is associated with prefrontal-leading beta-band coherence and Pop-out with parietal-leading gamma-band coherence: distinct frequencies for distinct systems (Buschman & Miller, 2007). Yet this account has faced mounting empirical and theoretical challenges (Awh et al., 2012; Katsuki & Constantinidis, 2014). Search difficulty varies continuously with target-distractor similarity and distractor-distractor similarity, rather than falling into two discrete categories (Duncan & Humphreys, 1992, 1989). Even within a single feature dimension such as orientation, search efficiency varies continuously with the heterogeneity of the distractors and with how targets and distractors are categorized, rather than switching between two modes (Wolfe et al., 1992). Optimal Bayesian observers with a single probabilistic computation, invoking neither parallel preattention nor serial inspection, reproduce the full range of search behavior (Eckstein, 2011, 2017). Most critically, intracranial recordings from human patients reveal that the majority of task-active sites are equally recruited in both Search and Pop-out across frontal, parietal, temporal, and medial temporal cortex, a result incompatible with the circuit-segregation prediction (Ossandón et al., 2012; Slama et al., 2021). If the same regions are active in both conditions, circuit segregation cannot be the answer. But what is? Demonstrating shared anatomy is necessary but not sufficient: it establishes that the same hardware is used, but leaves open how that shared neuroanatomy implements starkly different behavioral modes. The answer, we show in this study, lies in the temporal dynamics, specifically, in the oscillatory mechanisms that control how fast a shared circuit operates.

To test this, we recorded intracranial EEG from 19 presurgical patients (1,150 electrodes spanning occipital, parietal, frontal, premotor, and medial temporal cortex) while they performed a Pop-out versus Search task. Presurgical patients implanted for clinical monitoring offer a combination of spatial resolution, temporal resolution, and anatomical breadth unavailable to any non-invasive method, precisely what is required to simultaneously map active circuits and characterize their millisecond-scale oscillatory dynamics (Parvizi & Kastner, 2018). We report four convergent findings. First, time-frequency analysis reveals the same oscillatory signatures in both conditions: broadband high-frequency activity (HFA; 70–150 Hz) increases and alpha/beta (∼8–30 Hz) suppression propagating through the same posterior-to-frontal hierarchy, with no condition-specific frequency bands. Second, dynamic time warping demonstrates that the Search neural trajectory is a temporally stretched version of the Pop-out trajectory: the same sequence of neural events unfolding at a slower rate, with the degree of stretching predicting individual reaction time differences. Third, alpha-band (∼8–10 Hz) inter-areal coherence synchronizes the same posterior network in both conditions, with directed alpha coupling shifting systematically between time windows in a manner consistent with temporal scaling of information flow. Fourth, alpha phase gates local HFA amplitude at the same preferred phase in both conditions, demonstrating that the oscillatory machinery is shared across tasks; only the duration over which it is engaged differs. Together, these findings indicate that the cognitive flexibility distinguishing our two search conditions arises not from switching between dedicated circuits, but from temporal scaling of a single alpha-coordinated architecture. In essence, the same neural hardware operates at different speeds.

### 2.1. No frequency specificity between Search and Pop-out

We studied visual search because its behavioral dissociations have often been dichotomized into Pop-out and Search, implying two independent mechanisms. What we found is that these two conditions share the same cortical circuit, which we name the “posterior search network”. We analyzed iEEG from 19 presurgical patients with pharmacoresistant epilepsy (1,150 electrodes post-preprocessing) who performed a visual search task adapted from Li et al. (Li et al., 2010) (Fig. 1A–B). Participants identified a target among four triangle stimuli defined by color and orientation. In the Search condition, the target matched distractors in color; in Pop-out, it differed. Participants performed the visual search task with the expected behavioral dissociation between conditions: Pop-out responses were faster and less variable than Search responses (Fig. 1C), confirming that the two conditions differed markedly in difficulty. In Search the average reaction times (RTs) ranged from 0.655–1.382 s across participants (*M*=1.086, *SD*=0.203), while in Pop-out they ranged from 0.447–1.149 s (*M*=0.771, *SD*=0.190), with significantly slower RTs for Search compared to Pop-out (*t*(18)=14.573, *p*<0.0001).

**Fig. 1.**
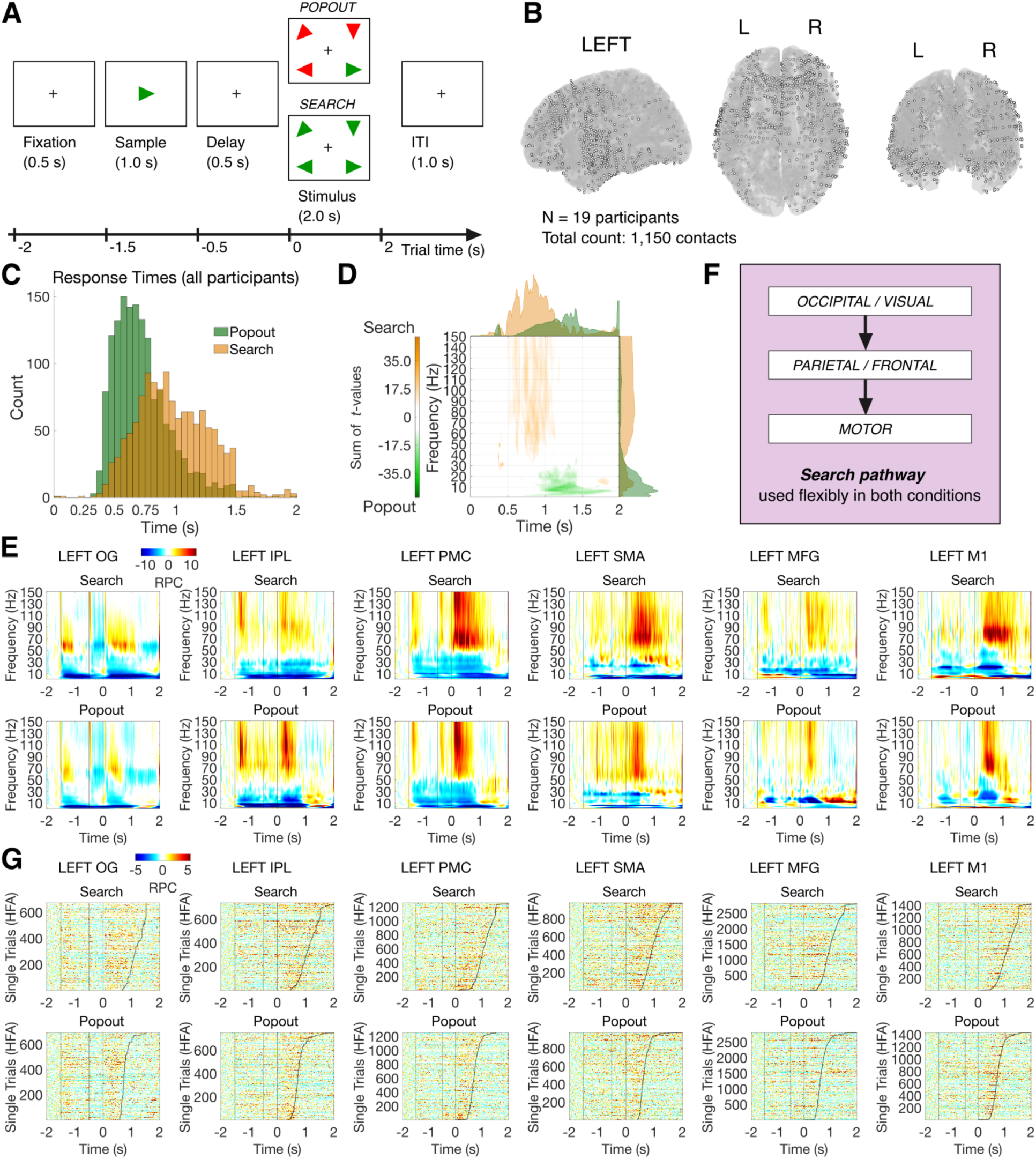
Task design, electrode coverage, and spectral dynamics of visual search in human iEEG. **A**, Trial structure. Each trial began with a fixation cross (0.5 s), followed by presentation of a sample triangle (1.0 s; target identity), a working memory delay (0.5 s), and the search display (up to 2.0 s). Time is shown relative to search display onset (t = 0). In the Pop-out condition, the target triangle was uniquely colored relative to the three distractors, rendering it immediately salient (an easy color-singleton search). In the Search condition, all triangles shared the same color, so the target had to be located by its orientation among heterogeneously oriented distractors (a difficult orientation search). The set size was four items in both conditions. The displays shown are illustrative examples; the orientations of the distractor triangles were randomized on every trial, and each distractor differed in orientation from the target. **B**, Intracranial electrode coverage across 19 presurgical patients (1,150 contacts total), shown on a standard brain template (left lateral, dorsal, and posterior views). **C**, Reaction time distributions pooled across all participants for Pop-out (green) and Search (orange), confirming faster and more compact responses in Pop-out. **D**, Time-resolved cluster-based permutation statistics comparing Search and Pop-out. Summed *t*-values are plotted as a function of time (relative to search display onset), with marginal frequency histograms. Positive (orange) and negative (green) clusters indicate significantly greater power in Search and Pop-out, respectively (*p_perm_*<0.05, 1,000 permutations). Condition differences span both high-frequency (Search > Pop-out) and low-frequency (Pop-out > Search) bands across a broad spectral range. **E**, Trial-averaged time-frequency power spectra (relative power change–RPC, z-scored to baseline) for Search (top) and Popout (bottom) across six left-hemisphere ROIs (OG, IPL, PMC, SMA, MFG, M1). Both conditions exhibit the same spectral signatures: broadband HFA increases (>70 Hz) time-locked to search display onset, accompanied by alpha/beta power decreases, with no evidence of condition-specific frequency specificity. **F**, Schematic of the search pathway identified from the spatiotemporal sequence of HFA activations in E: information flows from occipital/visual cortex to parietal/frontal cortex and onward to motor cortex. This hierarchy is engaged flexibly in both conditions, with closely matched dynamics at the great majority of sites. **G**, Single-trial HFA estimates (z-scored RPC) sorted by reaction time (black curve) for Search (top) and Pop-out (bottom) in each ROI. HFA increases are tightly time-locked to search display onset and terminate near the behavioral response in both conditions, confirming that the shared spectral dynamics observed at the trial-averaged level are present at the single-trial level. The shorter response window in Pop-out relative to Search reflects a difference in the duration of neural processing rather than its oscillatory character.

Participants were also more accurate in Pop-out than in Search. The accuracy was computed over responded trials after excluding time-out trials (trials with no response within the 2000 ms deadline). Accuracy was near-ceiling in Pop-out (*M*=97.05%, *SD*=2.67) and lower in Search (*M*=85.49%, *SD*=9.98; *t*(18)=-5.730, *p*<0.0001; Wilcoxon signed-rank *p*=0.00013). Equivalently, the error rate was low in Popout (*M*=2.95%, *SD*=2.67; 5 of 19 participants made no Pop-out errors) and higher, though still modest, in Search (*M*=14.52%, *SD*=9.98). Time-out trials were likewise less frequent in Pop-out (3.62% of trials) than in Search (16.86%; *t*(18)=4.181, *p*<0.001; Wilcoxon *p*=0.00065). This accuracy dissociation parallels the reaction-time dissociation, Pop-out is both faster and more accurate than Search (Supplementary Information, Section 1). Error trials and time-out trials were excluded from all neural analyses, which were therefore based on correct, responded trials only.

To determine whether Search and Pop-out recruit spectrally distinct neural processes, we performed a time-resolved cluster-based permutation test on the power spectra across all electrodes and ROIs (*p_perm_*<0.05 for the permutation test). Fig. 1D displays the significant between-condition difference. After stimulus onset, Search was associated with greater high-frequency power relative to Pop-out, while Popout was associated with greater low-frequency power relative to Search. These two effects reflect a single underlying difference in duration: the broadband high-frequency power increase is sustained for longer in Search (yielding the high-frequency Search higher than Pop-out clusters at late latencies), while the accompanying alpha/beta desynchronization is likewise more sustained in Search and recovers earlier in Pop-out (yielding, at later latencies, the low-frequency Pop-out higher than Search clusters). The significant clusters were not driven by any single dominant region, instead they were broadly distributed across both hemispheres and in both directions (19 ROIs for Search > Pop-out and 25 for Pop-out > Search; 12 ROIs showed effects in both directions; Supplementary Table 2, Supplementary Information, Section 2.1). Critically, the frequency distributions of the significant clusters revealed that significant differences were distributed broadly across the spectrum, without concentration in any particular, narrow frequency band selectively engaged by one condition. Rather than reflecting the selective recruitment of distinct oscillatory processes, these differences suggest that Search and Pop-out engage the same spectral architecture to different degrees, differing in the magnitude and duration of engagement, not in which frequencies are recruited.

This interpretation is supported by the trial-averaged time-frequency power spectra examined within-condition across six left-hemisphere ROIs spanning the dorsal attention network (OG, IPL, PMC, SMA, MFG, M1; Fig. 1E). In every region, both Search and Pop-out exhibited the same oscillatory signatures: a broadband HFA increase (>70 Hz) time-locked to search display onset, accompanied by a concurrent alpha/beta power decrease (∼8–30 Hz). These signatures were present in both conditions without exception. No region showed a condition-specific frequency band, i.e., no frequency range was selectively active in one condition but absent in the other. The primary differences between conditions were quantitative: Search was associated with sustained HFA increases of longer duration, consistent with the longer RTs, while Pop-out HFA responses were briefer and terminated earlier.

The spatiotemporal sequence of HFA activations across ROIs revealed a stereotyped cascade of activation common to both conditions: HFA increases emerged earliest in occipital and fusiform regions, propagated through parietal and frontal association cortices, and arrived last in premotor and primary motor cortex (Fig. 1F). This occipital → parietal/frontal → motor hierarchy constitutes a “search pathway” that was engaged flexibly in both Search and Pop-out, with closely matched cross-condition dynamics at the great majority of sites (see below), irrespective of whether target selection was driven by color or by orientation. The condition differences were not architectural (the same network was recruited in the same sequential order), but were instead temporal, reflecting differences in the duration over which each stage of the cascade remained active, and the RT differences between conditions.

Because this cascade was characterized at the level of pooled regions of interest, we additionally asked, at the level of individual contacts, whether pooling could obscure early condition differences at select ventral sites of the kind previously reported in fusiform cortex (Ossandón et al., 2012). A site-resolved analysis restricted to task-responsive occipito-temporal contacts confirmed that Search and Pop-out were statistically indistinguishable at the large majority of sites, while revealing a minority of fusiform contacts, including one with an early (<200 ms) onset, that showed stronger HFA in Pop-out than in Search. These effects were sparse and spatially restricted, consistent with transient bottom-up capture signals at select ventral sites rather than a distinct fast Pop-out circuit, and they do not alter the network-level conclusions (Supplementary Fig. 1, Supplementary Information, Section 2.2).

To verify that these trial-averaged observations were not an artifact of averaging across trials with heterogeneous dynamics, we examined single-trial HFA estimates sorted by reaction time (Fig. 1G). Across all six ROIs and in both conditions, HFA increases were time-locked to search display onset at the single-trial level, and the response in each trial terminated in close proximity to the behavioral response, as evidenced by the RT boundary curve tracking the offset of HFA activity. This trial-by-trial correspondence between neural dynamics and behavior was consistent across Search and Pop-out. Importantly, the shorter response window visible in Pop-out relative to Search directly mirrors the RT difference between conditions, confirming that the principal distinction between conditions is the duration of HFA engagement, not oscillatory character or spatial distribution.

To summarize, Fig. 1D shows between-condition differences, while Fig. 1E,G show the within-condition task-evoked responses. The near-absence of clusters in the early window (Fig. 1D) reflects the statistical equivalence of the early HFA responses across conditions (Fig. 1E,G). Positive (Search > Popout) clusters predominantly reflect broadband high-frequency activity sustained for longer in Search at late latencies, emerging only once the Pop-out responses terminate. Same for the negative (Pop-out > Search) clusters, reflecting a more sustained low-frequency desynchronization in Search (hence relatively greater low-frequency power in Pop-out, which recovers earlier). The late timing of the difference in Fig 1D is therefore expected, and is not in tension with the early, sustained within-condition HFA shown in Fig. 1E,G.

Together, these results establish that Search and Pop-out are not implemented by distinct neural oscillatory mechanisms. Instead, both conditions draw on a shared spectral repertoire (broadband activation and low-frequency suppression) deployed sequentially across a common cortical hierarchy, with condition differences arising from the timescale of processing rather than from the identity of the recruited circuits. What distinguishes Search from Pop-out is not which computations are performed, but how long the neural computation is sustained: a difference in temporal scale, not neural architecture.

These results contradict the circuit-segregation account. If Search and Pop-out recruit distinct oscillatory systems, as the classical parallel-versus-serial account would predict, we should observe condition-specific frequency bands (e.g., gamma for Pop-out, beta for Search, as reported in non-human primates) (Buschman & Miller, 2007) or region-specific recruitment patterns (e.g., ventral attention network for Pop-out, dorsal attention network for Search) (Corbetta & Shulman, 2002). We observe neither. Instead, both conditions engage the same spectral repertoire (HFA increases, alpha/beta suppression), deployed across the same cortical cascade (occipital → parietal/frontal → motor), with differences confined to the duration of engagement rather than the anatomy of recruited processes. This raises a new question: if the neural computations are the same, what determines how long they must run?

### 2.2. Dynamic time warping reveals temporal scaling underlies condition differences in visual search

The neural trajectory of Search is a time-stretched version of Pop-out. When we aligned the two conditions using dynamic time warping (DTW), a method that optimally maps one time series onto another by allowing flexible temporal rescaling (Sakoe & Chiba, 1978; Tavenard et al., 2020; Vintsyuk, 1968), the HFA signals from the same cortical regions became indistinguishable. What appeared to be two different neural responses was, after temporal alignment, a single response unfolding at two different speeds. This result is the neural signature of a shared circuit operating at different timescales.

We illustrate this with a representative example. In the left premotor cortex (Left PMC; Fig. 2A– B), the original Search and Pop-out HFA signals were visibly offset in time but otherwise structurally similar. Following dynamic time warping alignment, the two signals overlapped (Fig. 2A, warped signals), confirming that their temporal structures were equivalent. The warping path itself was linear across the search interval (Fig. 2B, top), indicating that the temporal scaling (dilation in Search or compression in Pop-out) is not confined to a single processing stage but reflects a uniform stretching of the entire post-stimulus neural trajectory, consistent with a change in processing speed rather than the insertion of an additional computational step. The warping offset curve, defined as the difference between the Pop-out and Search warping paths, was flat and not significantly different from zero during the pre-stimulus segment (Segment 1: *p*=0.987), confirming the absence of spurious pre-task offsets and validating this ROI for further analysis. During the task segment, the offset rose significantly above zero (Segment 2: *p*=0.0008), indicating that Pop-out and Search neural trajectories diverged in time, with Pop-out progressing faster along the shared trajectory than Search (Fig. 2B, bottom). Not because it performed a different computation, but because it performed the same computation more rapidly.

**Fig. 2.**
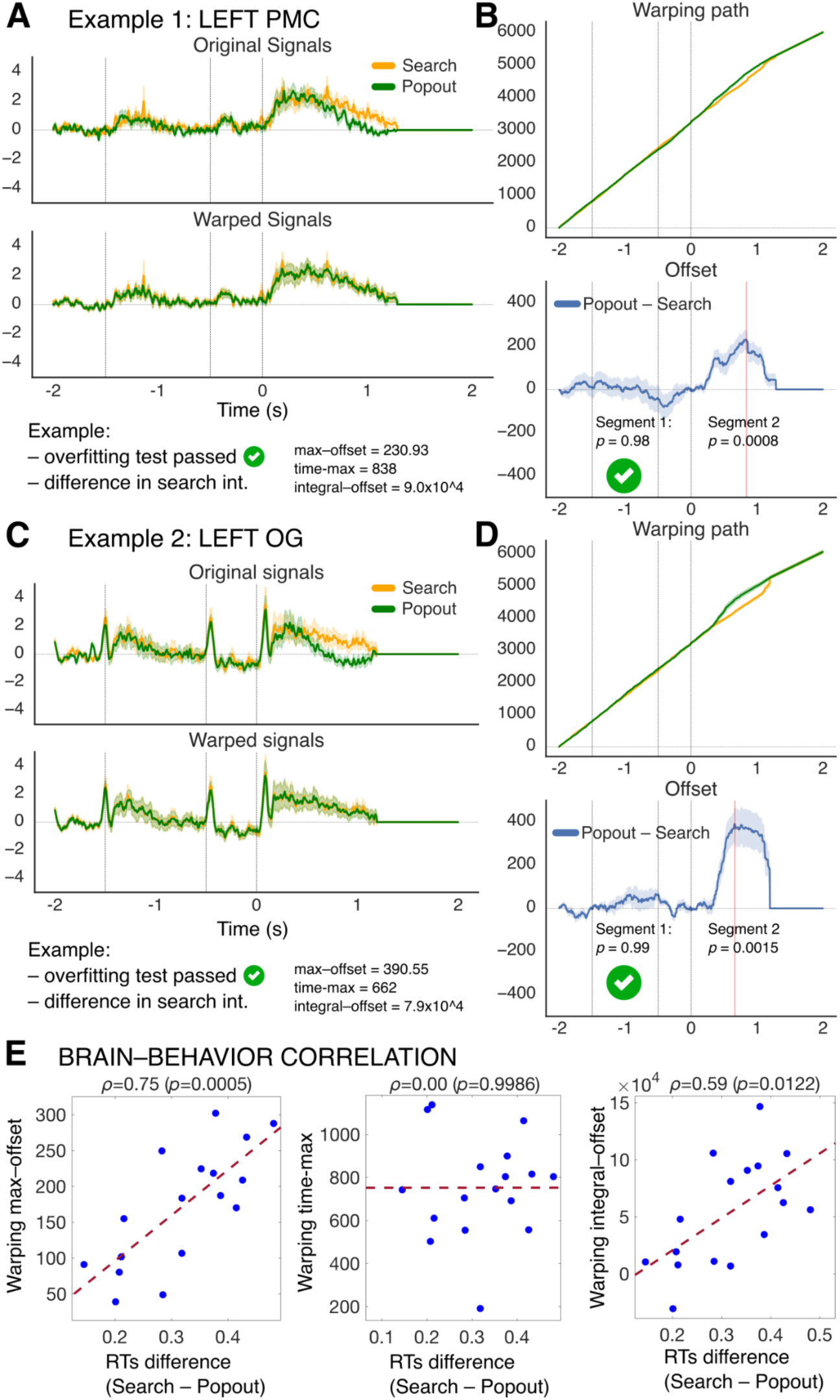
Dynamic time warping reveals temporal scaling as the mechanism distinguishing Search from Pop-out. **A**, Example of a valid region of interest (Left PMC). Trial-averaged HFA signals (z-scored; top), shown for Search (orange) and Pop-out (green), are structurally similar but temporally offset, with Search unfolding slower. After dynamic time warping (DTW) alignment (warped signals; bottom), the two traces overlap, demonstrating that the two conditions share a common underlying neural trajectory. In the ’Warped signals’ panel, the Pop-out trace (green) is re-indexed onto the Search time base according to the warping path, while Search (orange) is displayed unwarped. **B**, Warping path (top) and warping offset curve (Popout -Search; bottom) for the example in A. The warping path diverges after search display onset (t = 0), reflecting the temporal stretch of Search relative to Pop-out. The offset curve is not significantly different from zero during the pre-stimulus control segment (Segment 1: *p*=0.98, one-sided chi-squared test of variance), confirming the absence of spurious pre-task offsets, and rises significantly above zero during the task segment (Segment 2: *p*=0.0008, one-sided one-sample *t*-test). Shaded band indicates 95% confidence interval. This ROI passed validity criteria (max-offset = 230.93 ms; time-to-max = 838 ms; integral-offset = 9.0×10⁴). **C**, Same as panel A, showing another example of a region of interest (Left OG) that passed both validity criteria (Segment 1: *p*=0.99; Segment 2: *p*=0.0015). **D**, Warping path (top) and warping offset curve (Pop-out - Search; bottom) for the example shown in C (max-offset = 390.55 ms; time-to-max = 662 ms; integral-offset = 7.9×10⁴). **E**, Brain–behavior correlations. Per-participant peak warping offset (left; *ρ*=0.75, *p*=0.0005) and integral warping offset (right; *ρ*=0.59, *p*=0.0122) correlate positively with individual RT differences (Search - Pop-out), directly linking the magnitude of neural temporal dilation to behavioral slowing in Search (or otherwise temporal compression to behavioral acceleration in Pop-out). Time-to-maximum warping offset did not correlate with the RT difference (center; *ρ*=0.00, *p*=0.9986). Each blue dot represents one participant; dashed red lines show the best-fit linear regression. Eleven ROIs across both hemispheres passed both validity criteria and contributed to the participant-level analysis (left hemisphere: M1, MCC, MFG, OFC, OG, PMC, S1, SMA; right hemisphere: ACC, MFG, TPJ).

We illustrate a second example, the left occipital cortex (Left OG; Fig. 2C–D), whose warping offset appears more concentrated in the later processing period and might, from this single example alone, suggest stage-specific modulation, in which early sensory stages are minimally stretched and later stages involving binding and selection are stretched more. Similar to the Left premotor cortex, Left OG passed both validity criteria (Segment 1: *p*=0.999; Segment 2: *p*=0.0015). These two examples, one seemingly uniform and one seemingly non-uniform, illustrate precisely why the geometry of the warping path must be quantified across regions rather than read off individual offset curves, particularly because dynamic time warping pins the warping path to its endpoints and therefore produces a mid-interval offset hump under both uniform and stage-specific scaling. We address this next.

Across all recorded regions, eleven ROIs passed both validity criteria (absence of significant pre-stimulus offset and a significantly positive warping offset during the task interval): Left M1, Left MCC, Left MFG, Left OFC, Left OG, Left PMC, Left S1, Left SMA, Right ACC, Right MFG, and Right TPJ. Since the dynamic time warping analysis is designed to find an optimal temporal mapping between two signals, a significant positive warping offset by itself simply demonstrates such remapping, but not necessarily the presence of a shared trajectory between neural signals. To address this we derived measures of pre-warp and post-warp similarity, similarity change, and alignment cost in the post-stimulus interval, and compared them to a null model obtained by using mismatched ROI pairs (e.g., Search in Left OG vs. Pop-out in Left SFG–superior frontal gyrus, and so on). The results of this control analysis showed that the empirical Search and Pop-out signals become significantly more similar after warping, and at lower alignment cost, than what dynamic time warping produces for mismatched signals (Supplementary Fig. 2 and Supplementary Table 3, Supplementary Information, Section 3.1), demonstrating the presence of a genuine, shared neural trajectory between Search and Pop-out in each of the eleven ROIs that passed both validity criteria. These regions span the search pathway identified in the spectral analyses (Fig. 1F), from occipital and temporal cortex through parietal and premotor regions to frontal cortex, including both left-hemisphere dorsal attention network nodes and right-hemisphere frontal and parietal regions. The convergence of valid dynamic time warping results across this distributed network indicates that temporal scaling is a network-wide phenomenon, not restricted to any single processing stage. Every node of the search cascade slows down together.

Quantifying warping-path geometry across all valid ROIs shows that the temporal scaling is predominantly uniform. Because a mid-segment offset peak is an inevitable consequence of the endpoint constraints of dynamic time warping and is therefore not diagnostic of stage-specific stretching, we quantified features that discriminate uniform from non-uniform scaling: the linear-fit R² of each warping function, the trend in its instantaneous local slope across the task segment, and the skewness of the offset hump (Methods; Supplementary Information, Section 3.2). Across all eleven valid ROIs, the task-segment warping function was well approximated by a straight line (mean linear R² = 0.920, ranging between 0.857 in Left OG to 0.955 in Left MCC), the local slope showed only a weak, uniformly small positive trend across the segment (0.010–0.073 across ROIs), and the offset humps were close to symmetric, with skewness small and inconsistent in sign, ranging between -0.291 to +0.130 across ROIs (Supplementary Fig. 3 and Supplementary Table 4, Supplementary Information, Section 3.2). This pattern indicates approximately uniform stretching of the whole post-stimulus trajectory, with at most a mild and spatially inconsistent tendency toward greater stretching later in the segment. Notably, Left OG, whose single-example offset curve appeared late-concentrated (Fig. 2D), had the lowest linear R² among the valid regions, yet its warping function was still well fit by a straight line and its offset hump was essentially symmetric (skewness = -0.028) rather than late-skewed; its apparent non-uniformity therefore does not survive quantification. Thus the temporal scaling distinguishing Search from Pop-out is best described as a uniform dilation of the entire neural trajectory rather than a stage-specific stretch confined to late processing.

Critically, the observed temporal scaling was not a fixed property of the task. Instead, it varied parametrically across participants in direct proportion to their behavioral slowing. Participants whose neural trajectories stretched more showed larger Search-versus-Pop-out RT differences (peak warping offset: *ρ*=0.75, *p*=0.0005; Fig. 2E, left), in contrast to the time-to-maximum warping offset which did not correlate with the RT difference (Fig. 2E, center; *ρ*=0.00, *p*=0.9986). The cumulative temporal displacement over the entire task interval (integral warping offset) similarly correlated with the RT difference between conditions (*ρ*=0.59, *p*=0.0122; Fig. 2E, right), providing converging support for this brain–behavior relationship. This correspondence links the magnitude of neural temporal scaling to behavioral slowing across participants. Because the between-condition warping offset is itself indexed by trial duration (see RT-matched control, below), this correlation should not be read as independent evidence that neural scaling causes the RT difference; the evidence that the scaling is a genuine, response-speed–linked property of the circuit, rather than a re-description of duration, comes from the within-condition Fast-vs-Slow analysis (see below). This brain–behavior relationship demonstrates that individual differences in search efficiency are not due to recruiting different brain regions or oscillatory processes, but rather to how rapidly the same computation unfolds. Participants who are slower at Search relative to Pop-out traverse the same neural trajectory at a slower pace.

A response-bounded alternative could in principle explain the between-condition offset without any condition-specific scaling. If HFA is a fixed, response-bounded envelope, Search’s later responses would lengthen its trace and dynamic time warping would recover that duration. We tested this alternative directly against the data, using three controls: matching duration across conditions (below), testing signal specificity across frequency bands (see below and Supplementary Information, Section 3.5), and testing for shared internal structure against a mismatched-ROI null (see above and Supplementary Information, Section 3.1). An RT-matched control confirms that the between-condition difference is one of duration. We tested directly whether Search and Pop-out trials with matched reaction times share the same temporal profile. Within each participant we constructed RT-matched Pop-out and Search subsets by histogram stratification (50-ms bins; retaining the minimum number of trials between Search and Pop-out per bin), matching before pooling across participants (all but one participant retained ≥16 trials/condition; matched distributions statistically indistinguishable, all KS *p*>0.98, |SMD|<0.06) (Supplementary Fig. 4, Supplementary Information, Section 3.3). The trial-averaged HFA amplitude profiles of RT-matched Popout and Search overlapped closely in every ROI (Supplementary Fig. 5, Supplementary Information, Section 3.3), as predicted if the two conditions traverse a single shared trajectory. Applying the identical dynamic time warping pipeline to the RT-matched Pop-out and Search subsets, the task-segment warping offset collapsed toward zero (Supplementary Fig. 4, Supplementary Information, Section 3.3) and none of the eleven ROIs retained a significant positive task-segment offset once RT was equated (all *p*>0.15) (Supplementary Table 5, Supplementary Information, Section 3.3). Further, the warping offset did not exceed a within-participant condition-label-shuffle null (pooled peak-offset *p*=0.820 and integral *p*=0.490) in any of the ROIs (detailed results are provided in Supplementary Table 6, Supplementary Information, Section 3.3). Thus the between-condition temporal scaling is indexed by trial duration: when Search and Pop-out are matched on RT, no residual condition-specific trajectory difference remains. This is the expected signature of a single circuit whose only between-condition difference is the speed at which it runs, and it is complemented by the within-condition analysis below.

A within-condition analysis comparing Fast versus Slow trials establishes that the scaling is a genuine, response-speed–indexed property of the circuit. If the temporal scaling reflects processing speed rather than a Search-versus-Pop-out distinction, the same relationship should hold within a single condition between faster and slower trials. Within each task condition and participant, we split trials by a median split on RTs into Fast (shorter RTs) and Slow (longer RTs) subsets and applied the identical dynamic time warping pipeline, with the Slow series as reference so that a positive task-segment offset indicates that Slow is a temporally stretched version of Fast (all participants retained, subset sizes 22–32). The Fast and Slow HFA amplitude trajectories did not differ in the early post-stimulus window (0–300 ms) in any ROI valid in the original analysis, for either Search or Pop-out, mirroring the early between-condition equivalence and indicating that the two subsets differ in the temporal extent of the response rather than in early amplitude. Dynamic time warping revealed a positive task-segment offset in both conditions, in five of the eleven ROIs that were valid in the original analysis, with one or two additional ROIs trending to significance for each condition (Supplementary Table 7, Supplementary Information, Section 3.4). The warping offset exceeded a within-condition Fast/Slow-label-shuffle null (Supplementary Table 8 and Supplementary Fig. 6, Supplementary Information, Section 3.4): at the pooled level, the task-segment peak offset and integral both exceeded the null in both Search (peak offset 127.70 vs. null mean 17.44, *p*<0.001; integral 60,260.49 vs. null mean 178.27, *p*<0.001) and Pop-out (peak offset 92.43 vs. null mean 12.49, *p*<0.001; integral 23,578.78 vs. null mean -480.72, *p*<0.001). Regionally, the peak warping offset exceeded the null in 7 of 11 ROIs in Search and 8 of 11 in Pop-out, with 6 ROIs robust in both conditions (Left M1, Left MFG, Left OG, Left PMC, Left SMA, and Right TPJ) (Supplementary Fig. 6, Supplementary Information, Section 3.4). The integral metric exceeded the null in 5 of 11 ROIs in Search and 6 in Pop-out (robustly in Left M1, Left MFG, Left PMC, and Left SMA) (Supplementary Table 8, Supplementary Information, Section 3.4). Thus slower trials trace the same neural trajectory as faster trials, stretched in time, within each condition. We note that, in these within-condition analyses, while the significance of the alignment is less consistent across the eleven ROIs valid in the original analysis (possibly due to the reduction of trials per condition), the pooled result is unambiguous in both conditions and across offset metrics. Further, the ROIs showing a robust temporal scaling also showed a robust task-evoked HFA response (i.e., Left M1, MFG, OG, PMC, and SMA). Together with the RT-matched control (above), this indicates that a single cortical mechanism generates the search response at a rate indexed by reaction time, whether that rate varies between conditions or between trials within a condition.

To test whether the time warping results reflect a property of high-frequency activity (HFA) specifically, we repeated the dynamic time warping analysis, replacing HFA power with power in the beta-band (14–30 Hz) or in the alpha-band (8–13 Hz). We found that the effect was strongly selective to HFA. Of the 11 ROIs that met both validity criteria for HFA, only one, the left occipital cortex (Left OG), survived for both beta-band and alpha-band power. No additional regions reached joint significance in either control bands (Supplementary Table 9; Supplementary Information, Section 3.5). The single surviving ROI in both control bands is the primary visual area, where low-frequency (alpha/beta) power is itself strongly stimulus-driven. A residual temporal scaling signature there is therefore expected and, if anything, reinforces that the effect tracks task-relevant activity rather than mere elapsed time. We emphasize that beta-band power is not an inert control in our data, which shows robust task-related alpha/beta suppression (Fig. 1E). The fact that this task-modulated band nonetheless shows no network-wide temporal scaling argues specifically against a generic task- or response-duration account. If warping is merely a consequence of the longer stimulus-to-response interval in Search, beta suppression (equally bounded by that interval) should exhibit the same scaling, which it does not. A similar rationale applies to the alpha-band. These controls confirm that the dynamic time warping effects are specific for HFA, rather reflecting a general feature of any neural signal unfolding between search-display onset and the response.

A note on hemispheric lateralization. The connectivity and phase-coupling analyses that follow are restricted to the left hemisphere, where dynamic time warping yielded its most spatially extensive and statistically tractable results. This left-lateralization is unlikely to reflect sampling bias, as electrode coverage was in fact slightly denser on the right (612 contacts) than on the left hemisphere (538 contacts). Two non-mutually-exclusive factors more plausibly account for it. First, several participants with the densest and highest-quality posterior coverage (e.g., IR45, the only patient with Left OG electrodes) happened to be left-implanted, providing greater statistical power to resolve the underlying dynamics in this hemisphere. Second, participants responded with their right hand, which may have biased the search-to-response cascade toward left-hemisphere motor and premotor circuits. Importantly, the dynamic time warping analysis itself yielded valid temporal scaling effects in both hemispheres (eight left-hemisphere and three right-hemisphere ROIs), indicating that temporal scaling is not intrinsically left-lateralized. Future work using cohorts with denser bilateral coverage, or with left-handed responders, can dissociate these contributions directly.

If the brain is running the same computation at different speeds, what coordinates this neural computation? The answer lies in alpha-band oscillatory dynamics.

### 2.3. Alpha oscillations serve as the primary inter-areal coordination signal in the posterior search network

Having established that Search and Pop-out differ in the timescale rather than the architecture of neural processing (Figs. 1–2), we next asked: what neural mechanism controls this timescale? A strong candidate is oscillatory synchronization, which has been proposed to flexibly coordinate information flow across distributed cortical networks by rhythmically aligning the excitability of neuronal populations (Fries, 2005, 2015). If temporal scaling is implemented via oscillations, we should observe (1) a dominant frequency band that couples the regions of the search pathway, and (2) condition-specific modulation of coupling strength or directionality reflecting the faster (Pop-out) versus slower (Search) traverse of the shared trajectory. We computed inter-areal coherence and directed Granger-Geweke causality (GGC) (Geweke, 1982; Pagnotta et al., 2018a, 2018b) across 11 predefined left-hemisphere ROIs spanning the dorsal attention network. This set comprised the eight left-hemisphere ROIs that passed dynamic time warping validity criteria (M1, MCC, MFG, OFC, OG, PMC, S1, SMA; see previous section), supplemented by three posterior regions (IPL, FFG, MTL) included on a priori anatomical grounds as core nodes of the dorsal attention network and the posterior visual hierarchy. These additions ensured that the connectivity analyses captured the full posterior-to-anterior extent of the search cascade identified in Fig. 1F, regardless of whether each individual ROI met the per-ROI dynamic time warping criterion.

Grand-average coherence, collapsed across all inter-ROI connections, revealed that alpha-band synchrony (∼8–10 Hz) is the dominant mode of inter-areal coupling in this network, present in both Search and Pop-out (Fig. 3A). Coherence peaked sharply in the alpha range and decayed steeply at higher frequencies, with comparable spectral profiles in both conditions. Examination of pairwise coherence across individual connections revealed that alpha-band coupling was not uniformly distributed across the network, but was spatially constrained to a posterior subnetwork, which we term the “alpha-coherence zone”, encompassing occipital (OG), parietal (IPL), somatomotor (S1/M1), premotor (PMC), and medial temporal (MTL) cortex (Fig. 3B). More anterior frontal regions (MCC, SMA, MFG, OFC) showed little to no sustained alpha-band inter-areal coherence. This spatial selectivity was evident in the pairwise time-frequency coherence plots: connections within the posterior subnetwork (e.g., OG–IPL, OG–MTL, IPL– MTL, IPL–PMC) showed robust low-frequency coherence in both conditions, while connections at the anterior boundary (e.g., MFG–MTL, PMC–MFG) showed no apparent coupling in any band (Fig. 3C). Critically, the coherence profiles did not differ significantly between Search and Pop-out at any examined connection (no cluster survived correction; the smallest *p*-value from the permutation test was *p_perm_*=0.108), consistent with alpha-band inter-areal synchrony being a shared feature of both conditions rather than a distinguishing feature of either.

**Fig. 3.**
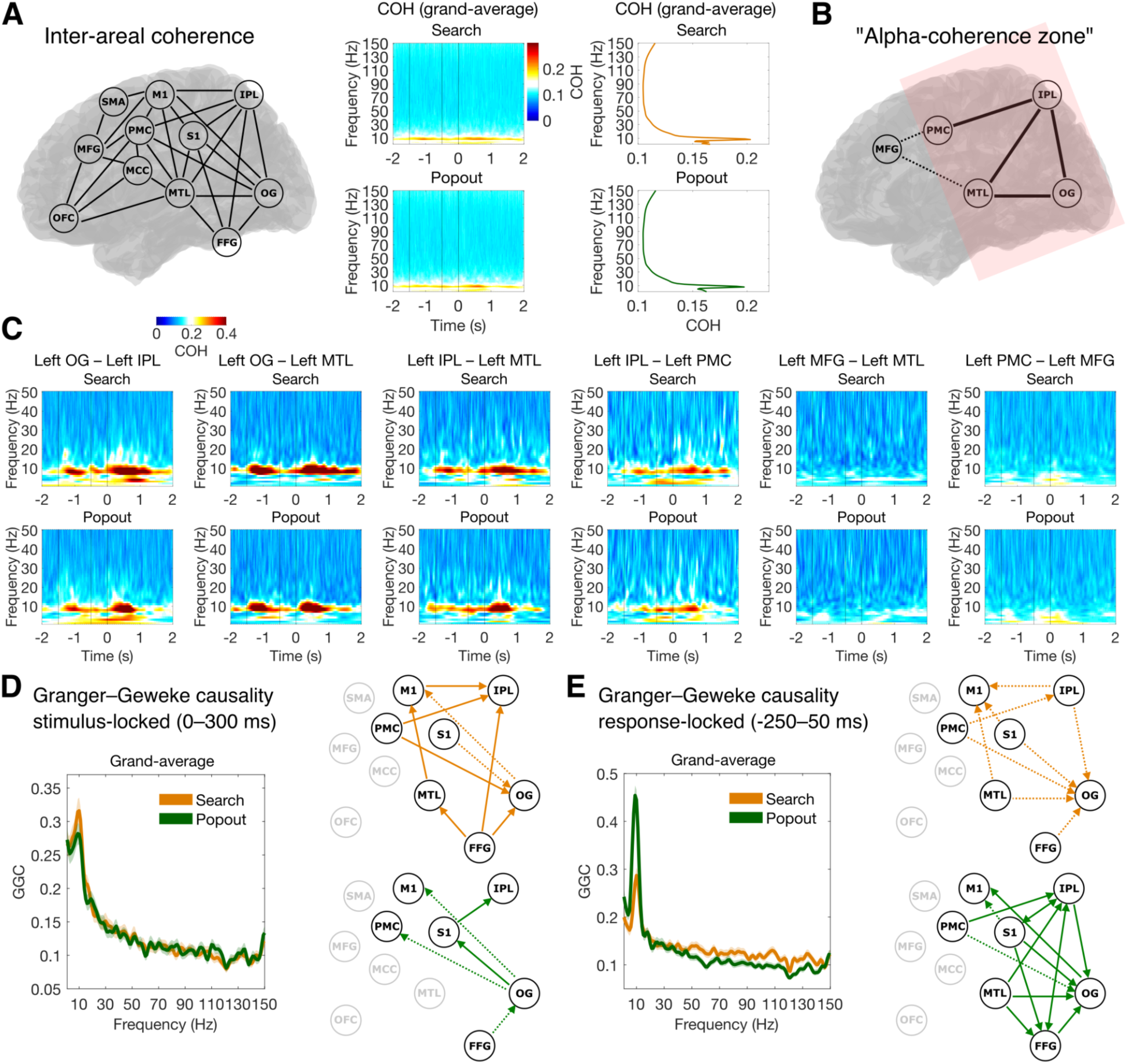
Inter-areal alpha-band coherence and directed connectivity are shared across Search and Pop-out but differ in their directed patterns. **A**, Grand-average inter-areal coherence (COH). Left: network of 11 predefined left-hemisphere ROIs over which pairwise coherence was estimated. Center: grand-average time-frequency coherence matrices, collapsed across all inter-ROI connections, for Search (top) and Pop-out (bottom). Right: corresponding time-collapsed coherence spectra for Search (orange) and Pop-out (green). In both conditions, coherence peaks sharply in the alpha band (∼8–10 Hz) and decays steeply at higher frequencies, establishing alpha-band synchrony as the dominant mode of inter-areal coupling shared across conditions. **B**, Anatomical extent of the “alpha-coherence zone.” Solid lines indicate ROI pairs with the strongest alpha-band coherence from the examples provided in C, forming a posterior subnetwork spanning occipital (OG), parietal (IPL), premotor (PMC), and medial temporal (MTL) cortex.

Despite this shared pattern of coherence, directed Granger-Geweke causality analysis revealed condition-specific differences in the pattern of alpha-band information flow that depended on the temporal window of analysis. In the early stimulus-locked window (0–300 ms), the grand-average Granger-Geweke causality spectrum peaked in the alpha band in both conditions, with Search showing marginally higher grand-average directed influence than Pop-out (Fig. 3D, left). At the level of individual connections, Search was characterized by a distributed pattern of significant directed alpha-band influence: feedforward connections spanned from occipital and fusiform cortex (OG, FFG) through parietal cortex (IPL) and onward to somatosensory and motor regions (S1, M1), with additional influence from MTL (Fig. 3D, right, top). Several of these connections showed stronger directed influence in Search than in Pop-out (solid arrows) after applying FDR correction: from FFG → OG (*p*<0.0001, *q*=0.0019), FFG → IPL (*p*<0.0001, *q*=0.0001), and FFG → MTL (*p*=0.0047, *q*=0.047), MTL → M1 → IPL (*p*=0.0007, *q*=0.012 and *p*=0.0005, *q*=0.010, respectively), as well as the directed connections from PMC → IPL (*p*<0.0001, *q*=0.0006) and PMC → OG (*p*<0.0001, *q*<0.0001). By contrast, Pop-out showed a sparser pattern of significant directed connections, including OG → S1 (*p*=0.0022, *q*=0.027) and S1 → IPL (*p*<0.0001, *q*<0.0001), predominantly confined to posterior regions (Fig. 3D, right, bottom), suggesting that the early feedforward propagation of alpha-band influence along the search pathway is more extensive in Search. The results revealed no detectable Granger-Geweke causality in the connections with anterior frontal ROIs (SMA, MCC, MFG, OFC), outlining a spatial boundary that is in line with the alpha-coherence zone identified before (Fig. 3)

In the peri-response window (−250 to +50 ms), this pattern reversed (Fig. 3E). Pop-out now showed a substantially denser and more widespread pattern of significant directed alpha-band connections than Search, spanning the full posterior-to-anterior extent of the network, from OG and FFG through IPL, S1, MTL, and M1 (Fig. 3E, right, bottom; on significant directed connections all *p*-values<0.0005, all *q*-values<0.0043). Search, by contrast, showed a more selective set of significant connections in this window, characterized by weaker strength than Pop-out (Fig. 3E, right, top). This temporal dissociation (broader early feedforward alpha coupling in Search, followed by broader peri-response alpha coupling in Pop-out) provides a directed-connectivity correlate of the temporal scaling effect identified by dynamic time warping: in Pop-out, the same network computations unfold faster, reaching the response-preparation stage earlier, shifting the window of maximal alpha-band directed influence toward the response epoch. This pattern is what the temporal scaling account predicts: if Search and Pop-out traverse the same computational stages in the same order, but at different speeds, then the timing of inter-areal alpha-band information flow should shift accordingly, not because the pattern of connectivity changes, but because the entire trajectory is stretched or compressed in time. The Granger-Geweke causality analysis provides independent convergent evidence for the dynamic time warping result at the network level.

Taken together, these results establish that alpha oscillations serve as the primary inter-areal coordination signal in the posterior search network. Alpha-band coherence is a shared, spatially selective feature of both conditions, while condition differences emerge in the directed patterns of alpha-band influence and their temporal evolution, consistent with a single shared circuit operating at different speeds.

Dashed lines indicate weaker or less consistent connections at the anterior boundary (MFG). More anterior frontal regions are absent, indicating that alpha-band inter-areal coherence is spatially constrained to posterior and medial temporal nodes of the dorsal attention network. Pink shading highlights this anatomical territory. **C**, Pairwise time-frequency coherence for six representative inter-ROI connections, shown for Search (top) and Pop-out (bottom) in each pair. Connections within the alpha-coherence zone (OG–IPL, OG–MTL, IPL–MTL, IPL–PMC) show sustained low-frequency coherence in both conditions; connections at the anterior boundary (MFG–MTL, PMC–MFG) show weaker coupling. The similarity between conditions across all pairs confirms that alpha-band inter-areal coherence is a shared property of both Search and Pop-out. **D**, Nonparametric pairwise Granger-Geweke causality (GGC) estimated in the early stimulus-locked window (0–300 ms). Left: grand-average GGC for Search (orange) and Pop-out (green), both peaking in the alpha band. Right: network diagrams of significant directed alpha-band connections for Search (top) and Pop-out (bottom). Search is characterized by a broadly distributed feedforward pattern of directed alpha influence, spanning from occipital and fusiform cortex through parietal and motor regions. Pop-out shows a sparser pattern, predominantly confined to posterior connections. **E**, Granger-Geweke causality (GGC) estimated in the peri-response window (−250 to +50 ms relative to response). Left: grand-average GGC. Right: network diagrams of significant directed connections. The condition asymmetry reverses relative to D: Pop-out now shows a denser pattern of directed alpha-band connectivity spanning the full posterior-to-anterior extent of the network, while Search shows a more selective pattern. This temporal dissociation (broader feedforward alpha coupling in Search during early processing, and broader alpha connectivity in Pop-out approaching the response) is consistent with the temporal scaling account, whereby the same network computations unfold at different speeds across the two conditions.

### 2.4. Alpha oscillations mediate these processes via phase-amplitude coupling with high-frequency activity

Having established that alpha oscillations serve as the dominant inter-areal coordination signal in the posterior search network (Fig. 3), we next asked how alpha mechanistically organizes local cortical processing. We tested for phase-amplitude coupling (PAC) between alpha phase and high-frequency activity (HFA), the putative signature of local neuronal activity (Lei et al., 2026; Leonard et al., 2023; Leszczyński et al., 2020; Ray & Maunsell, 2011). The temporal scaling account makes a specific and falsifiable prediction: if Search and Pop-out implement the same computation at different speeds, using a common oscillatory machinery, then the preferred alpha phase at which HFA is gated should be the same in both conditions. Only the duration over which this coupling is sustained should differ. We tested this prediction directly.

#### 2.4.1. Alpha–HFA phase-amplitude coupling is present in both conditions

The preferred alpha phase at peak HFA was the same in Search and Pop-out. Across all significant channels in all posterior ROIs, phase preferences clustered at the same angular location in both conditions (Fig. 4). If Search and Pop-out used distinct oscillatory mechanisms the preferred gating phase should differ between conditions, reflecting different functional relationships between alpha rhythm and local computation. It did not. The same alpha phase modulates local processing in both conditions. What changes between conditions is not the gate, but how long it stays open.

**Fig. 4.**
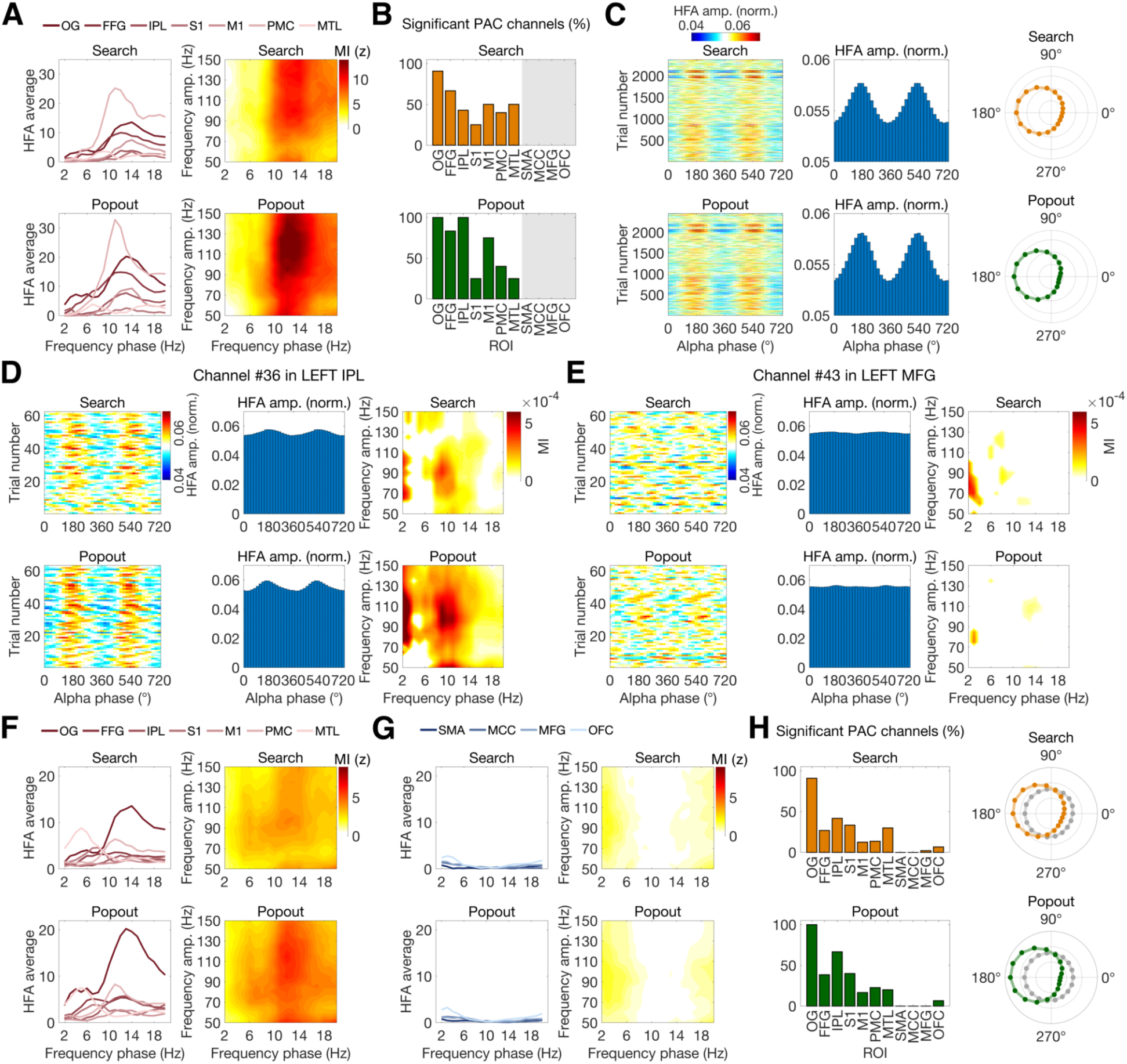
Alpha–HFA phase-amplitude coupling is a spatially selective, condition-invariant feature of the posterior search network. **A–C**, Single-participant data (patient *IR45*; the only participant with Left OG coverage). **A**, Left: mean HFA amplitude as a function of low-frequency phase for each ROI (color-coded lines; see legend), for Search (top) and Pop-out (bottom). Posterior regions, particularly OG, show prominent sinusoidal modulation of HFA amplitude as a function of alpha phase. Right: grand-average frequency-frequency modulation index (MI, z-scored against surrogate distributions) comodulograms, showing coupling between phase frequencies (x-axis, 2–20 Hz) and amplitude frequencies (y-axis, 50–150 Hz). A prominent alpha–HFA coupling peak (∼8–12 Hz phase, 70–150 Hz amplitude) is present in both conditions. **B**, Percentage of electrodes with statistically significant PAC (*z*>2.33, *p*<0.01, surrogate-based threshold) per ROI, for Search (orange) and Pop-out (green). Significant PAC channels are most prevalent in posterior regions (OG, FFG, IPL), in both conditions. **C**, Single-trial alpha–HFA PAC in the ROIs shown in B. Left: HFA amplitude as a function of alpha phase for individual trials (y-axis: trial number). Center: trial-and-channel-averaged phase histogram (bar plots), showing mean HFA amplitude as a function of alpha-band phase (0°–720°, two full cycles shown for clarity). Right: preferred alpha phase for each significant channel (polar plots). HFA amplitude is consistently maximal at a preferred alpha phase in both Search (orange) and Pop-out (green), with phase preferences clustering tightly across channels and trials. **D–E**, Single-participant data (patient *CP19*; a participant with coverage in Left IPL and Left MFG). **D**, Single-channel example of robust alpha–HFA PAC (channel #36, Left IPL). Left: single-trial HFA amplitude as a function of alpha phase. Center: trial-averaged phase histogram. Right: raw MI comodulogram. All three representations converge on a focal, consistent alpha–HFA coupling signature in both Search and Pop-out. **E**, Single-channel example of absent alpha–HFA PAC (channel #43, Left MFG). Same format as D. No systematic phase-dependent HFA modulation is observed in either condition, and the comodulogram shows no focal coupling. Together, D and E illustrate the spatial specificity of alpha– HFA PAC at the single-channel level: robust in the posterior cortex, absent in the prefrontal cortex. **F**–**H**, Group-level data (all participants with electrode coverage). **F**, Same format as A for posterior ROIs (OG, FFG, IPL, S1, M1, PMC, MTL). Alpha–HFA PAC is robust at the group level in both conditions, with OG showing the largest phase-dependent HFA modulation. **G**, Same format as A and F for anterior frontal ROIs (SMA, MCC, MFG, OFC). HFA amplitude shows no phase-dependent modulation and comodulograms are uniformly near-zero, confirming the spatial specificity of alpha–HFA PAC to posterior cortex. **H**, Group-level percentage of significant PAC channels per ROI (bar plots) and distribution of preferred alpha phases across all significant channels (polar plots), for Search (orange) and Pop-out (green). Significant PAC channels are most prevalent in posterior regions (OG, FFG, IPL) and decline steeply toward anterior frontal cortex (MCC, MFG, OFC), in both conditions. Phase preferences cluster consistently across participants and conditions.

We first established this in patient IR45, the only participant with electrode coverage in left occipital gyrus (OG), which is the region that showed the strongest alpha-band coherence and the largest dynamic time warping offset, providing a critical test case. In this patient, HFA amplitude was modulated as a function of alpha phase across multiple posterior ROIs, with OG showing the strongest modulation (Fig. 4A). The frequency-frequency modulation index (MI) comodulogram confirmed that this modulation was concentrated in the alpha–HFA frequency combination (∼8–12 Hz phase, 70–150 Hz amplitude), forming a focal coupling peak in both Search and Pop-out (Fig. 4A, right). The proportion of electrodes with statistically significant PAC (z>2.33, *p*<0.01, surrogate-based threshold) was highest in posterior regions in both conditions, peaking in OG (Pop-out: 100%; Search: 91%), FFG (83%; 67%), and IPL (100%; 43%), and declining through M1, MTL, PMC, and S1 (Fig. 4B). Although the absolute proportion of significant PAC channels was descriptively higher in Pop-out than Search across most posterior regions (not formally compared between conditions, for the reasons detailed in the Methods), the preferred coupling phase itself did not differ between conditions (see below).

Examination of single-trial PAC estimates revealed that HFA amplitude was maximal at a preferred alpha phase across individual trials in both Search and Pop-out, with no systematic difference in preferred phase between conditions (Fig. 4C). The phase histogram revealed a prominent sinusoidal structure, visible across two full alpha cycles (Fig. 4C, center). This sinusoidal structure, with HFA rising and falling predictably with the phase of every alpha cycle, means that local neural activity is not continuous, but discretized into rhythmic windows of excitability set by the alpha oscillation. The polar distributions of the preferred alpha phase across significant channels clustered tightly at the same angular location in both conditions (Fig. 4C, right), confirming that phase-gating architecture is not only present in both Search and Pop-out, but is locked to the same alpha phase regardless of which of the two tasks is performed. The coupling network, its mechanism, and its preferred alpha phase are shared across conditions. What differs is the duration over which coupling is sustained and, as a consequence, the prevalence of detectable coupling across channels, which is greater in Pop-out, consistent with the temporal concentration of coupling in the faster condition.

As a control, we asked whether this coupling is specific to the search period. Repeating the PAC analysis in the pre-stimulus interval ([-2000, 0 ms] relative to search-display onset, spanning target encoding and maintenance) revealed alpha–HFA coupling at the same preferred phase in both conditions (Supplementary Fig. 7; Supplementary Information, Section 4), indicating that alpha-phase gating of HFA is a general property of the posterior cortex engaged across task phases, rather than a process evoked specifically by visual search.

#### 2.4.2. Alpha–HFA PAC is spatially specific to the posterior search network

This coupling was sharply bounded by the spatial extent of the alpha-coherence zone identified before (Fig. 3). The comodulograms for anterior frontal ROIs (SMA, MCC, MFG, OFC) were uniformly near-zero across the entire frequency-frequency space, with no focal alpha–HFA peak in either condition. The transition between alpha-coherence zone ROIs with robust PAC and anterior frontal ROIs with no alpha– HFA PAC was sharp rather than gradual, defining a functional boundary that aligns precisely with the network identified in the connectivity analyses (Fig. 3).

This dissociation is visible at the level of single electrodes within the same participant. In patient CP19, a channel in Left IPL (channel #36) showed classic alpha–HFA coupling: single-trial HFA amplitude was systematically modulated by alpha phase, with a consistent preferred phase visible across all trials in both Search and Pop-out (Fig. 4D). The trial-averaged phase histogram confirmed a bimodal distribution, reflecting one HFA peak at each alpha cycle (Fig. 4D, center; two full cycles shown for clarity), and the raw MI comodulogram showed a focal hot spot precisely at the alpha–HFA frequency combination (Fig. 4D, right). This channel exemplifies the coupling profile observed throughout the alpha-coherence zone: strong, consistent, condition-invariant alpha–HFA PAC. A channel in Left MFG (channel #43), recorded simultaneously in the same patient during the same task, showed the opposite pattern (Fig. 4E). Single-trial HFA amplitude was uniformly distributed across alpha phases in both conditions, the phase histogram was flat with no preferred phase, and the MI comodulogram was diffuse with no focal coupling peak. This demonstrates the spatial boundary of alpha–HFA PAC within the alpha-coherence zone: even within the same participant and the same task, PAC is present in the posterior cortex and absent in the prefrontal cortex.

These single-participant findings generalized to the full cohort. Across all participants with electrode coverage, the group-level HFA amplitude modulation by alpha phase was present in posterior ROIs (OG, FFG, IPL, S1, PMC, MTL) in both conditions (Fig. 4F). In stark contrast, anterior frontal ROIs (SMA, MCC, MFG, OFC) showed no detectable phase-dependent HFA modulation (Fig. 4G–H). This spatial boundary is not incidental: it defines the functional architecture of the posterior search network, the alpha-coherence zone. Alpha–HFA coupling is restricted to the alpha-coherence zone identified before (Fig. 3), where alpha coherence was strongest and directed alpha influence was most prominent. Frontal regions participate in the search cascade (Fig. 1F) but are not governed by the same oscillatory mechanism. They may instead receive the output of alpha-coordinated posterior processing rather than implementing alpha-gated computations themselves.

This anatomical dissociation suggests a two-stage computational architecture. The posterior cortex implements alpha-coordinated sensory and association processing, which outputs to alpha-independent decision or motor preparation processes in the frontal cortex. The absence of PAC in frontal regions does not imply they are uninvolved. Indeed, HFA increases are robust in some of these areas (e.g., MFG and SMA; see Fig. 1E). Rather, this indicates that frontal processing operates outside the alpha-gated temporal framework that governs posterior computations. This distinction may reflect a functional hierarchy: posterior regions flexibly coordinate feature binding and spatial selection on an alpha timescale, while frontal regions integrate the output of this process to form categorical decisions or motor plans, a computation that may require different oscillatory dynamics (e.g., beta-band communication with motor cortex) (Engel & Fries, 2010) or non-oscillatory accumulation mechanisms (e.g., ramping activity) (Gold & Shadlen, 2007).

Taken together, these results establish that alpha–HFA phase-amplitude coupling is a robust, spatially selective, and condition-invariant feature of the posterior search network. Alpha oscillations coordinate inter-areal communication through phase synchrony and also organize local cortical processing by gating HFA activity to specific phases of the alpha cycle, in both Search and Pop-out, and across the same regions that showed the strongest alpha-band coherence and the most reliable dynamic time warping offsets. The invariance of preferred coupling phase across conditions further supports the shared-circuit account: the oscillatory architecture that controls local computations is not reconfigured between Search and Pop-out. In Search, it simply runs longer.

### 2.5. A unified oscillatory architecture for visual search

Across four complementary analyses, we show that the difference between our two search conditions is implemented by temporal scaling of a single, shared neural architecture. Search and Pop-out recruit the same cortical cascade with the same spectral dynamics (Fig. 1), differing primarily in how long the shared trajectory is sustained (Fig. 2). Alpha oscillations coordinate this process at two levels: synchronizing activity across the posterior search network and shifting the timing of directed inter-areal influence in a manner consistent with temporal compression (Pop-out) versus dilation (Search) (Fig. 3), and phase-gating local HFA within posterior cortex at a fixed preferred phase in both conditions (Fig. 4). Together, these findings demonstrate that, across these two conditions, flexibility is achieved not by switching between dedicated circuits, but by controlling the speed at which the same computation unfolds.

## 3. Discussion

The brain achieves cognitive flexibility in the two search tasks studied here not by selecting among dedicated circuits, but through reconfiguration of its temporal and oscillatory coordination, adjusting the speed at which a single circuit operates. This is the central finding of the present study, demonstrated here with the spatiotemporal precision of human intracranial EEG. This finding challenges a foundational assumption of systems neuroscience: that different behaviors require different neural architectures. Visual search is one of the most robust and widely replicated phenomena in cognitive psychology. Using visual search as a testing ground, we show that two processing modes long attributed to distinct neural systems are instead implemented by the same cortical cascade, the same oscillatory frequencies, and the same alpha-gated coupling architecture, differing primarily in how long that shared trajectory is sustained. This claim is supported directly by an RT-matched control: when Search and Pop-out trials are matched on reaction time within participants, their high-frequency activity profiles overlap and the between-condition warping offset is abolished, confirming that the two conditions differ in the rate at which a shared trajectory unfolds rather than in the computation it performs. This duration dependence is specific to high-frequency activity and is not a generic consequence of the longer Search interval: beta-band suppression, an equally response-bounded and robustly task-modulated signal in the same regions, shows no comparable temporal scaling, excluding an account in which any response-bounded signal would yield the same warping offset. The same temporal scaling is present within each condition, between faster and slower trials: slower trials trace a time-stretched version of the faster-trial trajectory in both Search and Pop-out. These results establish that temporal scaling is a general property of the circuit indexed by response speed, rather than a feature specific to the Search–Pop-out contrast. The oscillatory mechanism coordinating it (alpha phase-gating of high-frequency activity at the same preferred phase across conditions) is indistinguishable between modes. This suggests that, for contrasts of this kind, the more informative question may not be “which system” a given behavior recruits, but “at what speed” a shared system operates.

In the domain of visual search, this conclusion resolves a tension that has persisted for over four decades. Treisman and Gelade’s Feature Integration Theory (A. M. Treisman & Gelade, 1980) proposed that feature search operates via parallel preattentive processing while conjunction search requires serial attention-dependent binding. This dichotomy was subsequently mapped onto distinct neural systems, distinct frequency bands of inter-areal coherence, and distinct cortical networks (Buschman & Miller, 2007; Corbetta & Shulman, 2002). Other work challenged this dichotomy, progressively weakening the circuit-segregation account: Duncan and Humphreys showed that search efficiency falls on a continuum rather than a categorical boundary (Duncan & Humphreys, 1989); Wolfe’s Guided Search model proposed that all search involves both parallel priority computation and serial selection (Wolfe, 2021); Eckstein demonstrated that optimal Bayesian observers, invoking neither parallel nor serial mechanisms, account for the full range of search behavior (Eckstein, 2011). Our results provide a mechanistic account for this alternative model: Search and Pop-out traverse the same neural pathway in the same order, coordinated by the same oscillatory architecture, at different speeds. The distinction Treisman drew between “serial” and “parallel” search may therefore reflect, at the neural level, the same computation running at different rates. We emphasize that our design does not test this identification directly. Both of our conditions are feature searches, and a decisive test would require conditions that differ in set size and in whether the target is defined by a conjunction of features. What Wolfe’s Guided Search correctly identified as shared computational steps are implemented by a single occipital-to-frontal cascade. What Eckstein’s Bayesian framework correctly identified as a single underlying computation is realized neurally through the sequential activation of this cascade, with alpha coordinating network synchrony and local neural computation.

“Serial search” in this temporal scaling account does not mean spotlight-like spatial scanning (classical Treisman’s FIT), but serial progression through processing stages, with progression speed determined by salience and guidance strength. We should be explicit about the grain of description at which this claim is made. Our measures are trial-averaged, region-level envelopes of high-frequency activity. They resolve the sequence and duration of stages in a distributed cortical cascade, not the individual comparisons by which any single item is accepted or rejected. What we claim is that this cascade, and the alpha coordination accompanying it, is traversed at a rate indexed by response speed, and that no stage or oscillatory mechanism appears at the network level in one condition that is absent in the other. We do not claim that every elementary operation within a stage is itself slowed. Quantification of the warping-path geometry indicates that this scaling is predominantly uniform: the neural trajectory is dilated approximately evenly across the post-stimulus interval rather than stretched selectively at a single late stage. We note that the mid-interval peak of the warping-offset curve is not evidence of stage-specific stretching, since dynamic time warping produces such a peak under uniform scaling as well. This uniform, whole-trajectory dilation is consistent with a single global change in processing rate, the same computation running slower, rather than the insertion or selective slowing of one processing stage; it also argues against the simplest bounded-accumulation account, in which non-decision (sensory and motor) time is held fixed and only a decision stage is prolonged. Separately, the directed alpha-connectivity analysis shows a condition-dependent shift in the timing of inter-areal coupling, from broader early feedforward coupling in Search to denser peri-response coupling in Pop-out, consistent with faster arrival at response preparation in Pop-out; this concerns the temporal organization of network coordination and is not, by itself, a claim about non-uniform stretching of the local HFA trajectory.

This leaves open a question that our data do not settle. A trajectory that takes longer may do so because the same operations each proceed more slowly, or because the same operations, proceeding at the same rate, are executed more times, as an account in which distractors are rejected iteratively would require. At the level of trial-averaged regional activity, these are not distinguishable, and our data do not localize where in the trajectory any additional operations would be inserted. What can be said is that both produce a longer traversal of the same circuit, and neither introduces a stage or a coupling regime absent from the other condition. Our phase-amplitude coupling results suggest what the natural unit of such iteration would be: high-frequency activity is modulated at the same preferred alpha phase in both conditions, which makes the alpha cycle a candidate quantum of processing. We did not count cycles, however, and this remains an interpretation of the coupling result rather than a measurement of it. The distinction is testable in principle: if difficulty is paid for in cycles rather than in rate, the number of alpha cycles between display onset and response should scale with the number of items to be rejected, which a design that varies set size could establish and ours cannot. We emphasize that both readings are versions of the claim we do make. Whether the circuit runs slower or runs more times, it is the same circuit, coordinated in the same way, and behavioral difficulty is expressed as the duration of its engagement.

Because both of our conditions can be described as a single decision process run at two rates, it is worth stating what our data do and do not settle about such an account. Three observations constrain it. The temporal scaling is distributed across the whole cascade rather than confined to one stage: significant warping offsets were obtained in early visual cortex (Left OG), in somatosensory and motor cortices (Left S1, Left M1), and in premotor and supplementary motor cortex (Left PMC, Left SMA), and the within-condition Fast-versus-Slow analysis recovers the effect over a comparable span (Left OG, MFG, PMC, SMA, and M1). The scaling is specific to high-frequency activity (HFA), since beta-band suppression is bounded by the same stimulus-to-response interval yet shows no network-wide dilation. Further, it is present within a single condition, where no condition comparison is involved. Together these argue against a model that holds sensory and motor time fixed and prolongs only a decision stage. They do not, however, adjudicate among accumulator models more generally. A model in which drift rate and non-decision time both vary between conditions is fully consistent with our results, and is in effect the temporal-scaling account expressed in accumulator vocabulary, since it holds that the entire processing chain rather than one stage of it runs at a different rate. Nor do our data license a mapping of individual regions onto the components of such a model: sustained occipital high-frequency activity during search may reflect ongoing evidence sampling rather than sensory encoding alone. What our results establish is the neural form the speed difference takes, a single cortical cascade coordinated by a single oscillatory mechanism and traversed at a rate indexed by response speed, rather than the algorithmic description best suited to summarizing it.

The broader implication is that cognitive flexibility may not require diverse anatomy (Buzsáki & Draguhn, 2004; Fries, 2015). The posterior search network identified here, consisting of occipital, fusiform, parietal, and premotor cortex coupled by alpha synchrony, is architecturally stable across conditions. What changes is temporal dynamics: how long each stage is sustained, when directed influences are exerted, and at what rate local computations proceed. This suggests a general principle: the brain achieves cognitive flexibility through temporal scaling of fixed circuits; the same hardware supports different modes by operating at different speeds. This may generalize beyond visual search, with the potential to reframe debates in cognitive control, working memory, and decision-making, where dual-process accounts have been proposed. Rather than asking which system is engaged, we might instead ask how fast a shared system operates and what oscillatory mechanisms coordinate the shared local-processing trajectory.

Mechanistically, alpha-band oscillations support temporal scaling through a dual coordination mechanism: inter-areal synchronization of posterior cortical regions and local phase-gating of high-frequency activity within those regions. This goes beyond prior characterizations of alpha as primarily an inhibitory signal (Jensen & Mazaheri, 2010; Klimesch, 2012; Pagnotta et al., 2020, 2022) or a general correlate of attentional engagement (Worden et al., 2000). The alpha coherence network spans occipital, parietal, premotor, and medial temporal cortex, precisely the regions showing the strongest HFA modulation and the most reliable temporal scaling; while directed alpha-band coupling shifts systematically between early feedforward (Search-dominant) and peri-response (Pop-out-dominant) windows in a manner consistent with temporal scaling of information flow. Crucially, alpha phase gates local HFA at the same preferred phase in both conditions (Fig. 4C, F), demonstrating that the oscillatory machinery is shared across tasks; only the duration over which it operates differs. This phase-gating mechanism provides a concrete implementation of how alpha organizes the timing of local computations: HFA, which indexes population activity, is modulated only during specific phases of the alpha cycle, effectively discretizing continuous processing into rhythmic temporal windows. If processing is organized into such windows, the alpha cycle becomes a natural unit in which the duration of a computation can be expressed, and a difference in difficulty can in principle be paid for either by lengthening each window or by traversing more of them. The greater prevalence of significant PAC (proportion of channels) in Pop-out does not reflect a broader or different circuit (the preferred phase and spatial boundary of coupling are identical across conditions), but follows from temporal scaling itself: compressing the same computation into fewer, less dispersed alpha cycles concentrates phase–amplitude coupling within the analysis window, whereas dilation in Search disperses it. Furthermore, the absence of network-wide temporal scaling in the alpha-band, as revealed by the control analysis using dynamic time warping, is consistent with the role our framework assigns to alpha. In our account, alpha does not itself constitute the scaled local-processing trajectory; rather, it coordinates that trajectory through inter-areal phase synchrony and phase-amplitude gating of HFA.

Our findings refine, rather than oppose, the influential non-human primate account of visual search proposed by Buschman and Miller (Buschman & Miller, 2007). They reported that prefrontal cortex signals target location before the parietal cortex during top-down conjunction search, while the parietal cortex leads during bottom-up feature search, with the two modes distinguished by different frequency bands of inter-areal coherence (beta for top-down Search, gamma for bottom-up Pop-out). However, the empirical foundation for the “frontal-leads-parietal” claim during Search is narrower than the study suggests. The cumulative-distribution analysis on which it rests (their Fig. 2D) shows LPFC and FEF reach significance at only ∼40 ms and ∼50 ms before saccade, respectively, while LIP reaches significance ∼32 ms after saccade, in the Search condition (Buschman & Miller, 2007). This means that the entire effect occupies a 70–80 ms window enclosing the saccade itself, and selectivity emerges roughly 220 ms after stimulus onset given their search reaction times. A signal that resolves peri-saccadically to post-saccadically is naturally interpreted as a saccade target-selection or motor-readout process rather than as the top-down control of search per se, and a search-control claim would be more directly tested in a stimulus-locked window. Notably, Buschman and Miller’s follow-up study reframed the relevant frontal signal as an FEF-localized serial spotlight clocked by ∼25 Hz oscillations, with no comparable ordering effect in dlPFC (Buschman & Miller, 2009), which is compatible with a temporal, oscillatory account of search. Our human iEEG data extend that trajectory: rather than two frequency-specific systems, we find a single alpha-coordinated posterior network whose rate of operation, not whose architecture, distinguishes fast from slow search. The residual disagreement between these two accounts plausibly reflects species differences and the absence in our paradigm of the strict eye-fixation requirement that engages FEF serial-spotlight dynamics in macaques.

Our results support and extend the fMRI-based dorsal/ventral attention network framework (Corbetta & Shulman, 2002). The alpha-coherence zone we identify (occipital, fusiform, inferior parietal, premotor, and medial-temporal cortex) overlaps with the canonical dorsal attention network: our IPL coverage spans portions of the intraparietal sulcus and supramarginal gyrus, and our PMC ROI is adjacent to and partially overlapping with the frontal eye fields. Our observation that these regions are equivalently recruited in both Search and Pop-out with largely overlapping dynamics aligns with recent human iEEG findings showing extensive overlap in task-active sites and activation dynamics across conditions (Ossandón et al., 2012; Slama et al., 2021). Our results specify the oscillatory mechanism and timing by which this network operates, revealing a critical functional boundary: alpha-mediated phase-amplitude coupling is spatially restricted to the alpha-coherence zone identified in our connectivity analyses (Fig. 3B), while anterior frontal regions (SMA, MCC, MFG, OFC) show robust HFA increases but no alpha–HFA coupling (Fig. 4E). This suggests a two-stage computational architecture in which posterior regions implement alpha-gated sensory and association processing, the output of which is passed to frontal regions that implement alpha-independent decision and motor preparation. Frontal cortex may operate via different oscillatory dynamics, such as beta-band communication with motor cortex during response preparation (Engel & Fries, 2010), or via non-oscillatory mechanisms such as ramping evidence accumulation (Gold & Shadlen, 2007). This parcellation of alpha-gated versus alpha-independent processing stages may explain why prior studies using whole-brain fMRI or scalp EEG, which average across functionally distinct regions, have produced mixed results regarding the role of alpha in visual search.

Several limitations warrant consideration. First, intracranial recordings provide unparalleled spatiotemporal resolution but necessarily involve sparse and clinically determined electrode placement (Parvizi & Kastner, 2018). Not all patients had coverage of all regions of interest, and the locations sampled were dictated by clinical need rather than experimental design. However, the fact that our core findings (shared spectral dynamics, temporal scaling, alpha coherence, and phase-amplitude coupling) replicate across patients with heterogeneous electrode placements and across multiple independent ROIs argues for generalizable features of the posterior search network, alpha-coherence zone, rather than artifacts of sampling bias. Second, our patient population consisted of individuals with pharmacoresistant epilepsy, raising the question of whether epilepsy or antiepileptic medications might alter oscillatory dynamics. We mitigated this concern by excluding electrodes in or near seizure onset zones and by confirming that behavioral performance (RT distributions) matched that of neurologically intact participants in prior studies. Nevertheless, replication in non-clinical populations using high-density scalp EEG, MEG source reconstruction, or invasive recordings in non-human primates performing analogous tasks would strengthen confidence in generalizability. Third, our findings are correlational rather than causal. While the tight coupling between neural temporal dilation/compression and behavioral slowing/acceleration, the systematic shifts in directed alpha connectivity, and the condition-invariant preferred phase for alpha–HFA coupling together provide strong mechanistic constraints, direct causal evidence requires interventional approaches. Fourth, our connectivity and phase-coupling analyses were restricted to the left hemisphere for the reasons detailed in the Results section, and although dynamic time warping yielded valid temporal scaling effects in both hemispheres, the bilateral nature of the alpha-coordinated posterior network remains to be confirmed in cohorts with denser right-hemisphere coverage and with left-handed responders. Fifth, because the task required a two-alternative (left/right) localization judgment, correct responses can in principle include lucky guesses, most notably in the less accurate Search condition. This does not affect our conclusions: neural analyses were restricted to correct, responded trials; any residual guessing biases the temporal-scaling and coupling estimates toward the null rather than generating them; the error rate was low in both conditions in the key left-occipital dataset (patient IR45); and the within-condition Fast-versus-Slow and reaction-time-matched analyses, which do not depend on the between-condition accuracy difference, reproduce the effect. Sixth, the set size was held constant at four items. Our design therefore does not measure the reaction-time–by–set-size function that distinguishes efficient from inefficient search, and we make no claim about the search mode of either condition. We characterize the two task conditions operationally, as an easy color-singleton search and a difficult orientation search, and the difference we report is one of difficulty and speed. For the same reason, our data cannot determine whether the slower condition reflects item-by-item inspection of the display or a single slower decision process, and we do not claim to resolve the parallel-versus-serial question. The question our design does address is whether the two conditions are implemented by distinct circuits or by one, and on this the reaction-time-matched control is directly informative: once reaction time was equated within participants, no residual condition-specific difference remained in any of the eleven regions tested. A study that varies set size, and that includes a conjunction condition, would allow the temporal-scaling account to be tested against search efficiency directly. Seventh, we did not control for perceptual grouping among the search items. In a display of four oriented triangles, subsets of items can group by orientation or by collinearity, and such grouping can produce emergent features that speed or slow responses, including the “false pop-out” effect in which a grouped subset captures attention without containing the target (Orsten-Hooge et al., 2015). Two features of the design bound, without eliminating, this possibility: i) the four item positions were fixed in the same four quadrants throughout the experiment and are therefore identical across conditions, and ii) distractor orientations were randomized on every trial, so any orientation-based grouping varied from trial to trial rather than acting as a fixed property of one condition. Nonetheless, we note that the Search displays were homogeneous in color and are for that reason more groupable overall than the Pop-out displays, in which the target is set apart by color. We regard this as one plausible contributor to the difficulty of the Search condition rather than as a confound for the question our design addresses, since the temporal-scaling account is indifferent to what makes a given trial slow: whatever its cause, a slower trial traverses the same trajectory more slowly, which is what the within-condition Fast-versus-Slow analysis shows directly.

These limitations point directly to testable predictions. If alpha oscillations causally implement temporal scaling, then rhythmic perturbation of alpha via transcranial alternating current stimulation (tACS) should parametrically modulate visual search reaction times. Specifically, alpha-frequency tACS applied to posterior parietal or occipital cortex should selectively affect the Search condition more than Pop-out, because Search requires sustained alpha-mediated coordination and is thus more sensitive to exogenous disruption of the alpha clock. Furthermore, the phase of alpha stimulation relative to the endogenous oscillation should matter: stimulation that aligns with the endogenous preferred phase for HFA gating should facilitate visual search, while anti-phase stimulation should impair it. Beyond visual search, if temporal multiplexing is a general principle, then other tasks exhibiting dual-process or parallel-versus-serial dissociations should show shared neural substrates with parametric differences in oscillatory timescale rather than categorical differences in regional recruitment. Automaticity in motor sequence learning, traditionally attributed to a shift from cortical to striatal control (Doyon et al., 2009), may instead reflect temporal compression of cortico-striatal circuits operating at different speeds. Implicit versus explicit memory retrieval, historically dissociated into hippocampal and striatal systems (Squire & Dede, 2015), may reflect flexible coordination of these regions over different timescales, with recent evidence suggesting these systems interact dynamically rather than operating in isolation (Packard & Goodman, 2013). Type 1 versus Type 2 reasoning, distinguished as automatic versus deliberate cognitive processes (Evans & Stanovich, 2013; Kahneman, 2011), may arise from the same neural architecture operating under different temporal constraints. This view is supported by critiques of strict dual-system accounts (De Houwer, 2019).

The present findings support a framework for understanding cognitive flexibility, in which the brain achieves behavioral versatility not by maintaining separate neural systems for different processing modes, but by controlling the speed at which a shared computation unfolds. We demonstrate this for two visual search tasks that differ in difficulty, implemented by a shared alpha-coordinated architecture. Whether it extends to search tasks that differ in kind rather than in difficulty, and to the other domains in which dual-process architectures have been invoked, is a prediction this work motivates. Testing it would shift the question, in those domains, from “which system?” to “at what speed?”.

## 4. Methods

### 4.1. Participants

Intracranial recordings were obtained from 19 human participants (age range 21–69 years, *M*=34.57, *SD*=13.95; 11/8 female-male ratio; 17 were right-handed and 2 ambidextrous), who underwent presurgical epilepsy evaluation at the California Pacific Medical Center, USA (N=6), the University of California Irvine Medical Center, USA (N=12), and Oslo University Hospital, Norway (N=1). All patients provided written informed consent to participate in this study, which was approved by the Institutional Review Boards of the participating hospitals, as well as the Committee for the Protection of Human Subjects at the University of California, Berkeley. In Norway, the study was also approved by the Regional Committees for Medical and Health Research Ethics. All patients had IQ above 85, normal or corrected-to-normal vision, and no known deficits in visual perception or color vision. Patients were tested when they were fully alert and willing to participate.

### 4.2. Stimuli, design, and procedures

The stimuli and task design were adapted from Slama et al. (Slama et al., 2021), which provides a full description of the complete cohort of 23 participants; the current study analyzed a subset of 19 of those patients. Stimuli consisted of acute isosceles triangles, either red or green, presented against a white background on a 15.6-inch Windows laptop LCD screen. Stimulus presentation and response collection were controlled using E-Prime 2.0 software (Psychology Software Tools). The laptop was positioned approximately 40–60 cm from the participant. Each triangle subtended 3.8–5.7° of visual angle at the base and 4.3–6.4° at the height. On the search display, each triangle was positioned 5.7–8.6° horizontally and 3.8–5.7° vertically from a central fixation cross, measured to the center of each stimulus.

Each trial began with a green fixation cross presented for 1000 ms (intertrial interval; ITI), the first 500 ms of which served as a pre-task baseline. The fixation cross then turned black for 500 ms (fixation interval), after which a single target triangle was displayed at the center of the screen for 1000 ms (sample interval). Participants were instructed to encode this target, defined by its color (red or green) and orientation (one of eight possible orientations: 0°, 45°, 90°, 135°, 180°, 225°, 270°, or 315°). Following a 500 ms working memory delay (working memory interval), the search display appeared, consisting of four triangles, including the target, arranged around the fixation cross (Fig. 1A). The positions of the four triangles (the target plus the three distractors) were located in the four quadrants of the screen (upper-left, lower-left, upper-right, and lower-right), and did not change during the course of the experiment. The target was randomly located in one of the quadrants (each trial included a target). The orientations of the distractor triangles were randomized for each trial (i.e., the configuration of the search display varied trial to trial), and the orientation of each distractor differed from that of the target. Participants indicated whether the target triangle was located in the left or right half of the display by pressing the left or right arrow key, or the left or right button of an external mouse, depending on the physical constraints of the recording environment. The trial terminated upon response, or after a maximum of 2000 ms if no response was made.

The experiment comprised 128 trials organized into four blocks of 32 trials each, with two conditions intermixed: Search and Pop-out. In the Search condition, all four triangles shared the same color (all red or all green), requiring participants to locate the target based on its orientation alone (a difficult orientation search). In the Pop-out condition, the target was uniquely colored relative to the three distractors (e.g., a red target among green distractors, or vice versa), rendering the target perceptually salient (an easy color-singleton search). Set size was fixed at four items in both conditions and was not manipulated. Half of the trials in the experiment were in the Search condition and half were in the Pop-out condition. Half of the targets were presented in the left visual field and half in the right. To synchronize neural and behavioral recordings, stimulus onset and offset signals were transmitted from the stimulus computer to the neural recording hardware via analog channels, using a photodiode at U.S. hospital sites and a TRS audio connector at Oslo University Hospital.

Behavioral performance was quantified for each participant and condition (Search, Pop-out). Single-trial responses were classified as correct, incorrect (error), or time-out (no response within the 2000 ms response deadline). Accuracy was defined as the proportion of correct responses among responded trials (correct / [correct + error]), excluding time-out trials, and the error rate as its complement (1 - accuracy). We additionally counted the number and proportion of time-out trials per condition. Condition differences in accuracy, error rate, and time-out trials were tested across participants with paired-samples *t*-tests, as for the reaction-time analysis. Because these measures are bounded and can approach ceiling or floor, we additionally report the non-parametric Wilcoxon signed-rank test for each comparison. Error trials and time-out trials were excluded from all subsequent neural analyses (time-frequency power, high-frequency activity, dynamic time warping, inter-areal connectivity, and phase-amplitude coupling), which were computed on correct, responded trials only.

### 4.3. iEEG data acquisition and preprocessing

All patients were implanted with electrocorticography (ECoG) or stereo EEG (sEEG) depth electrodes to determine the site of seizure onset for surgical resection. All electrodes implantations were guided entirely by clinical needs and considerations. Full details of data acquisition, anatomical reconstruction, electrode localization, and preprocessing are reported in Slama et al. (Slama et al., 2021). As described therein, power-line noise was attenuated during preprocessing by notch-filtering the line frequency and its harmonics; the large majority of patients (18 of 19) were recorded at North American sites with a 60 Hz line frequency. Briefly, iEEG was recorded across three clinical sites using Nihon Kohden and NicoletOne/ATLAS digital acquisition systems, at native sampling rates of 512, 1000, 1024, or 5000 Hz. The present study used a subset of 19 patients selected on the basis of acquisition sampling rates of 1000 Hz or integer multiples thereof (5000 Hz); data acquired at 5000 Hz were downsampled to 1000 Hz prior to analysis. We analyzed iEEG data from 1,150 electrodes. Electrode locations were determined by co-registering each patient’s post-implantation CT scan with their preoperative T1-weighted MRI using Freesurfer 5.3.0 and the FieldTrip toolbox, and electrodes were realigned to the preoperative cortical surface to correct for post-craniotomy tissue displacement. Anatomical labels were assigned by a neurologist (R.T.K.) based on inspection of fused MR/CT images in native space using BioImage Suite.

### 4.4. Power analysis

Time-frequency power spectra for each electrode were estimated using a Morlet wavelet transform (*ω₀*=6) over the interval [-3.0, +2.0] s relative to sample onset, across frequencies 1–150 Hz in 1 Hz steps. For each ROI, spectral estimates were pooled across all electrodes and smoothed with a moving average (window = 41 samples) prior to visualization. Condition differences (Search vs. Pop-out) were assessed using a nonparametric cluster-based permutation test, using a two-tailed dependent *t*-test (*df*=18, *p*<0.05, alpha level distributed over both tails by multiplying the *p*-values with a factor of 2 prior to thresholding), 1,000 permutations, and *p_perm_*<0.05 for the permutation test. Statistical testing was restricted to the post-stimulus interval [0, +2.0] s.

Single-trial high-frequency activity (HFA; 70–150 Hz) was estimated as a measure of local broadband cortical activity. The lower bound of the HFA band was set at 70 Hz to isolate broadband high-frequency activity, an index of local population spiking (Ray & Maunsell, 2011), from lower-frequency narrowband gamma oscillations, and to avoid the 50–70 Hz range most affected by the 60 Hz line-noise notch. Single-trial HFA was estimated using a Morlet wavelet transform (*ω₀*=6, with zero-padding), identical in implementation to the full time-frequency power analysis, but with the resulting per-frequency power estimates averaged across the HFA band (70–150 Hz) to yield a single broadband amplitude envelope per trial and electrode. As described in Section 4.2, only correct, responded trials entered this and all subsequent neural analyses; error and time-out trials were excluded.

### 4.5. Dynamic time warping

We used a dynamic time warping (DTW) analysis (Sakoe & Chiba, 1978; Vintsyuk, 1968) to quantify a hypothesized time-scaling effect between Pop-out and Search conditions, assessing the idea that the two conditions share the same neural trajectory but unfold at different speeds, with Search being a temporally stretched version of Pop-out. To test whether Search and Pop-out share a common neural trajectory unfolding at different timescales, we applied DTW (Tslearn) (Tavenard et al., 2020) to trial-averaged, z-scored time series derived from the spectrogram for the HFA frequency bands of interest (70–150 Hz), for stimulus-locked ([-2.0, +2.0] s) epochs. Z-scored time series were truncated at each participant’s mean Search RT to remove inter-individual RT confounds. For each electrode, DTW was applied separately to the pre- and post-event segments. For display purposes (Fig. 2A,C), the Search trace is shown unwarped and serves as the reference time base, and the Pop-out trace is resampled onto that time base according to the warping path. The transformation therefore rescales the Pop-out time axis, so displayed Pop-out amplitudes (’warped signals’) are not identical to the original ones. All statistical analyses were performed on the warping path and the derived warping offset, not on the resampled traces, which serve a purely illustrative purpose. The warping offset, which was defined as the difference between the Pop-out and Search warping paths, was used as a time-resolved index of temporal scaling. In the location-aggregated analysis, electrodes were pooled within each ROI across participants. ROI validity required two conditions: (i) absence of significant pre-stimulus offset, assessed via a one-sided chi-squared test of variance (null: variance≤300²), and (ii) a significantly positive warping offset during the task segment, assessed via a one-sided one-sample *t*-test against zero. Eleven ROIs met both criteria. In the participant-aggregated analysis, electrodes were pooled across these ROIs within each participant, and the peak warping offset during the task segment was correlated with each participant’s mean RT difference (Search - Pop-out) using Pearson correlation.

To confirm the presence of a shared trajectory between neural signals in Search and Pop-out, above and beyond the successful temporal remapping signaled by a significantly positive warping offset during the task interval, we performed a control analysis using a null based on mismatched ROI pairs. We divided the ROIs into two groups: the first group, named ’valid’ group, comprised the eleven regions that passed both validity criteria and showed significantly positive warping offset during post-stimulus interval (see above); the second group included all the other ROIs (’invalid’). We first performed a ’matched’ DTW approach on the empirical signals in Search and Pop-out in each valid ROI. We then performed a ’mismatched’ DTW approach, by taking the Search signals from the valid ROI, and warping them to the Pop-out traces randomly drawn from the invalid group. We compared matched vs. mismatched (Mann– Whitney U test, *p*<0.05), looking at four metrics: *pre-warp similarity* (correlation of Search and Pop-out signals in the post-stimulus task segment before DTW), *post-warp similarity* (same as the previous one, but after DTW), *similarity change* (post-minus pre-), and *alignment cost* (integral offset). We found that, in all valid ROIs, pre-warp and post-warp similarities were significantly higher in matched compared to mismatched (highest *p*=0.0350 in Right ACC and *p*=0.0112 in Right TPJ, for the two measures respectively), while the alignment cost was significantly lower (highest *p*=0.0208 in Left OG). The similarity change was larger for mismatched than the matched pairs in ten of the eleven ROIs (largest *p*=0.0407 in Left MCC), all valid ROIs except Right ACC (*p*=0.6134). These results, summarized in Supplementary Fig. 2 and Supplementary Table 3 (Supplementary Information, Section 3.1), demonstrate that the empirical Search and Pop-out traces become more similar and at lower cost, than what DTW could produce for unrelated or mismatched signals.

To distinguish uniform from stage-specific temporal scaling, we characterized the geometry of the task-segment warping function for each valid ROI. Because DTW pins the warping path to its endpoints, the warping offset is necessarily zero at both segment boundaries and forms a mid-segment hump under either hypothesis; the existence of a late-peaking offset is therefore not by itself diagnostic. For each contact we expressed the task-segment DTW correspondence as a warping function mapping Search time onto aligned Pop-out time, normalized to a common [0, 1] domain to accommodate per-contact truncation lengths, and computed (i) the coefficient of determination (R²) and residual RMSE of a straight-line fit (uniform scaling predicts R² near 1); (ii) the Pearson correlation of the instantaneous local slope with normalized time (a non-zero trend indicates time-varying, non-uniform scaling); and (iii) the skewness of the warping offset across the task window (a symmetric hump is consistent with uniform scaling, a late-skewed hump with stage-specific stretching). Metrics were computed per contact and averaged within ROI. The results are summarized in Supplementary Fig. 3 and Supplementary Table 4 (Supplementary Information, Section 3.2).

To test whether the temporal-scaling relationship between conditions reflects a general dependence of the neural trajectory on response speed, we performed two control analyses: i) RT-matched control (Supplementary Information, Section 3.3), ii) Within-condition Fast vs. Slow (Supplementary Information, Section 3.4). In the first analysis (RT-matched control), we repeated the DTW analysis on RT-matched Pop-out and Search subsets, to test whether the temporal scaling survives once trial duration is controlled. Within each participant, the two conditions’ RTs were pooled and partitioned into fixed 50-ms bins (25 ms as robustness check); within each bin we retained min(n_Pop-out, n_Search) trials per condition by seeded subsampling without replacement, discarding bins present in only one condition. Matching was performed within participants, pooling across participants only afterwards; participants retaining <16 trials/condition were excluded. Match quality was verified per participant (two-sample Kolmogorov–Smirnov test, standardized mean difference, overlap coefficient; retained if KS *p*>0.05 and |SMD|<0.1). Matched series were truncated at each participant’s mean RT over the pooled matched trials and passed unchanged through the DTW pipeline. Each task-segment metric (peak/integral offset) was compared, per ROI and pooled, against a within-participant condition-label-shuffle null (1,000 permutations; two-sided *p*).

In the second analysis (within-condition Fast vs. Slow), we divided each participant’s trials, separately within Search and within Pop-out, into Fast and Slow subsets by a rank-based median split on single-trial RT (equal halves; the single middle trial dropped when the count was odd), excluding participants retaining <16 trials per subset (all participants met this floor in both conditions). Per-trial baseline-z-scored HFA was averaged over each subset, and both series were truncated at the participant’s mean RT of the Slow subset (computed per condition). The Fast and Slow series were passed unchanged through the DTW pipeline with the Slow series as reference, so a positive task-segment offset indicates that Slow is a temporally stretched Fast. Before warping, early (0–300 ms) mean HFA was compared between Fast and Slow with a two-sided paired *t*-test across contacts per ROI. Each task-segment metric was compared, per ROI and pooled, against a within-condition Fast/Slow-label-shuffle null (1,000 permutations, truncation held at each participant’s real Slow-subset mean); because the hypothesis is that a genuine speed difference produces more alignment than chance relabeling, we report the one-sided upper-tail permutation *p*. Search and Pop-out were analyzed separately.

Finally, to assess the specificity of the observed temporal scaling effect for HFA, we repeated the DTW analysis and quantified the time-stretching effect between Pop-out and Search conditions for two control low-frequency bands: beta (14–30 Hz) and alpha (8–13 Hz). The only region that showed significant DTW effects for both beta-band and alpha-band power, in addition to HFA, was the Left OG (Supplementary Table 9, Supplementary Information, Section 3.5). No additional ROIs reached significance in either control band.

### 4.6. Connectivity: coherence

Time-varying phase-based connectivity was estimated between all pairs of electrodes spanning eleven predefined left-hemisphere ROIs, which were selected to combine the data-driven DTW-valid set (eight ROIs: M1, MCC, MFG, OFC, OG, PMC, S1, SMA) with three core anatomical nodes of the dorsal attention network and posterior visual hierarchy (IPL, FFG, MTL), included on a priori anatomical grounds. Using the same Morlet wavelet decomposition as for the power analysis (*ω₀*=6, zero-padding, 1–150 Hz at 1 Hz steps), coherence (COH) was computed from the trial-averaged complex wavelet coefficients for each electrode pair. Epochs were stimulus-locked ([-3.0, +2.0] s) and trial counts were equalized across conditions within each participant by random subsampling prior to estimation. Connectivity estimates were averaged across all electrode pairs contributing to each inter-ROI connection and grand-averaged across connections for visualization. Condition differences (Search vs. Pop-out) were assessed using a nonparametric cluster-based permutation test (two-tailed dependent *t*-test 1,000 permutations, *p_perm_*<0.05), analogous to the power analysis (described above).

### 4.7. Connectivity: Granger-Geweke causality

Directed functional connectivity was assessed using nonparametric pairwise Granger-Geweke causality (GGC) (Geweke, 1982). GGC was estimated between all electrode pairs spanning predefined inter-ROI connections, covering all left-hemisphere ROI pairs from the coherence analysis (see above). For each electrode pair, GGC was estimated within fixed time intervals extracted from stimulus-locked epochs in the [0, +300 ms] post-stimulus window, as well as a peri-response window [-250, +50 ms]. Spectral decomposition within each interval used a multitaper approach (time-bandwidth product *NW*=2) over 1– 500 Hz at 1 Hz resolution (Pagnotta et al., 2018a, 2018b). Following the approach of Pracar et al. (Pracar et al., 2026), GGC was estimated pairwise (bivariate) for each electrode pair, yielding directed influence in both directions and a net influence index (GGC₁→₂ - GGC₂→₁), normalized within each pair by the maximum GGC value across directions and conditions. Trial counts were equalized across conditions by random subsampling prior to estimation. Condition differences (Search vs. Pop-out) in directed and net GGC were assessed in the alpha band (9–10 Hz) using paired-samples *t*-tests, and net influence per condition was tested against zero using one-sample *t*-tests, with FDR correction for the number of ROI pairs tested using the Benjamini-Hochberg procedure (*q*<0.05). A grand-average was additionally computed by pooling all directed estimates across predefined left-hemisphere ROI pairs for visualization purposes.

### 4.8. Phase-amplitude coupling (PAC)

Phase-amplitude coupling (PAC) was quantified using Tort’s modulation index (MI) (Tort et al., 2010) computed within the stimulus-locked interval [0, +2000 ms] relative to search display onset, separately per electrode and condition (Search, Pop-out). The MI was estimated across a frequency-frequency grid spanning phase frequencies of 2–20 Hz (1 Hz steps, 2 Hz bandwidth) and amplitude frequencies of 30–150 Hz (5 Hz steps, bandwidth equal to twice the phase center frequency), using 18 phase bins. Epochs were extended by 500 ms on each side of the analysis window and trimmed after filtering to prevent edge artifacts. To standardize MI values and assess significance, following the approach of Daume et al. (Daume et al., 2024), 200 surrogate MI distributions were generated per channel by randomly shuffling the phase-amplitude pairing across trials. A normal distribution was fit to each surrogate distribution, and observed MI values were z-scored accordingly. A channel was considered to show significant PAC if its z-scored MI exceeded *z* = 2.33 (one-tailed, *p*<0.01). The proportion of channels with significant PAC was assessed per ROI and per condition. Because the two conditions differ in processing duration, and because Tort’s MI is sensitive to the number of oscillatory cycles in the analysis window, PAC was assessed within each condition separately (against channel-specific surrogates) rather than by a direct between-condition comparison of modulation-index magnitude: a fixed full-length window would confound coupling strength with the differing fraction of task-active time across conditions, whereas a shorter equal-length window would not afford stable modulation-index estimation. Between-condition differences are therefore reported as the proportion of channels with significant PAC per ROI, and interpreted descriptively. In a separate analysis, we further derived single-trial MI estimates to visualize trial-by-trial PAC dynamics and characterize alpha-band phase preferences in individual participants and channels. As a control, the identical PAC pipeline was additionally applied to the pre-stimulus interval [-2000, 0 ms] relative to search-display onset; full details and results are reported in Supplementary Information, Section 4.

## Acknowledgements

This work was supported by the National Institutes of Health (Grants R01NS021135, R.T.K.; Conte Center P50MH109429, R.T.K.; U19NS107609, R.T.K., J.J.L.). A.-K.S. and T.E. acknowledge support from the Research Council of Norway (Grant 240389/F20).

## Author Contributions

M.F.P. conceptualized the study. S.J.K.S. developed the experimental paradigm, supervised data collection, and contributed to data curation. D.K.-S., K.D.L., T.E., A.-K.S., and J.J.L. contributed to data collection. M.F.P. and H.Z. developed the analytical framework. M.F.P. and R.T.K. wrote the manuscript.

## Supplementary Information

### SI – 1. Behavioral analyses

Behavioral performance was quantified for all 19 participants. For each participant, single-trial responses were separated by condition (Pop-out, Search) and classified as correct, incorrect (error), or time-out. Trials on which no response was registered within the 2000 ms search-display window (time-out trials) were identified and excluded from the accuracy computation. Accuracy was defined as the proportion of correct responses among responded trials (correct responses divided by the sum of correct and error responses), and the error rate as its complement. In addition to accuracy and error rate, we counted the number and proportion of time-out trials per condition. To test whether performance differed between conditions, we compared Search and Pop-out across participants using paired-samples *t*-tests, as done for the reaction-time analysis reported in the main text. Because these measures are bounded and can approach ceiling (accuracy) or floor (errors, time-outs), we additionally report the non-parametric Wilcoxon signed-rank test for each comparison as a robustness check. Error trials and time-out trials were excluded from all neural analyses.

Participants performed the task accurately in both conditions, with a marked advantage for Popout. Accuracy was higher in Pop-out (*M*=97.05%, *SD*=2.67, range 91.67–100%) than in Search (*M*=85.49%, *SD*=9.98, range 62.90–98.25%), with a significant difference between the two (*t*(18)=-5.730, *p*<0.0001; Wilcoxon signed-rank *p*=0.00013). Equivalently, the error rate was significantly lower in Popout (*M*=2.95%, *SD*=2.67, range 0–8.33%; five of nineteen participants made no Pop-out errors) and higher in Search (*M*=14.52%, *SD*=9.98, range 1.75–37.10%). Time-out trials were likewise less frequent in Popout (*M*=2.32 trials, 3.62% of trials, *SD*=3.90) than in Search (*M*=10.79 trials, 16.86% of trials, *SD*=11.89; *t*(18)=4.181, *p*<0.001; Wilcoxon *p*=0.00065). Error and time-out trials were excluded from all following neural analyses.

Because the task required a two-alternative (left/right) localization judgment, chance performance was 50%, so errors raise the possibility that some nominally correct responses reflect lucky guesses. Under the worst-case assumption that every error is a random guess, an error rate *E* implies a guessing prevalence of approximately *2E* (half of guesses are correct by chance): about 6% of responded trials in Pop-out and about 29% in Search, with a corresponding fraction (*E*: ∼3% in Pop-out, ∼15% in Search) of nominally correct trials potentially reflecting lucky guesses. This is an upper bound, as many errors in a demanding orientation search are systematic (for example, mislocalization arising from erroneous orientation binding) rather than random 50/50 guesses, so the true guessing prevalence is lower. Three considerations indicate that this does not affect our conclusions. First, guessing is negligible in the dataset on which the mechanistic results most depend: patient *IR45*. This is the only participant with left occipital (OG) coverage, which is the region carrying the between-condition power difference, the sustained high-frequency activity, the dynamic-time-warping and phase-amplitude-coupling effects, and the strong alpha-band coherence and directed-connectivity differences. In patient *IR45*, the error rate was low in both conditions (3.17% in Popout, 5.66% in Search; Supplementary Table 1), corresponding to a worst-case guessing prevalence of only ∼6% and ∼11%. Second, all neural analyses were computed on correct, responded trials, and any lucky-guess trials retained in that set would, if anything, bias the results toward the null. A trial answered by a guess is one in which the target was not localized through the genuine search process, so its inclusion adds atypical, more variable single-trial trajectories that attenuate, rather than manufacture, the temporal-scaling and phase-amplitude-coupling effects we report. Third, the decisive tests of the temporal-scaling account do not rely on the between-condition accuracy difference: the within-condition Fast-versus-Slow analysis (Section 3.4, below) recovers the same temporal scaling within Pop-out alone, where errors are only ∼3%, and the reaction-time-matched control (Section 3.3) equates the conditions and collapses the between-condition offset. Neither result can be explained by guessing. We therefore report the Search error rate transparently while concluding that guess-related contamination does not undermine the single-circuit, speed-scaling interpretation.

**Supplementary Table 1.**
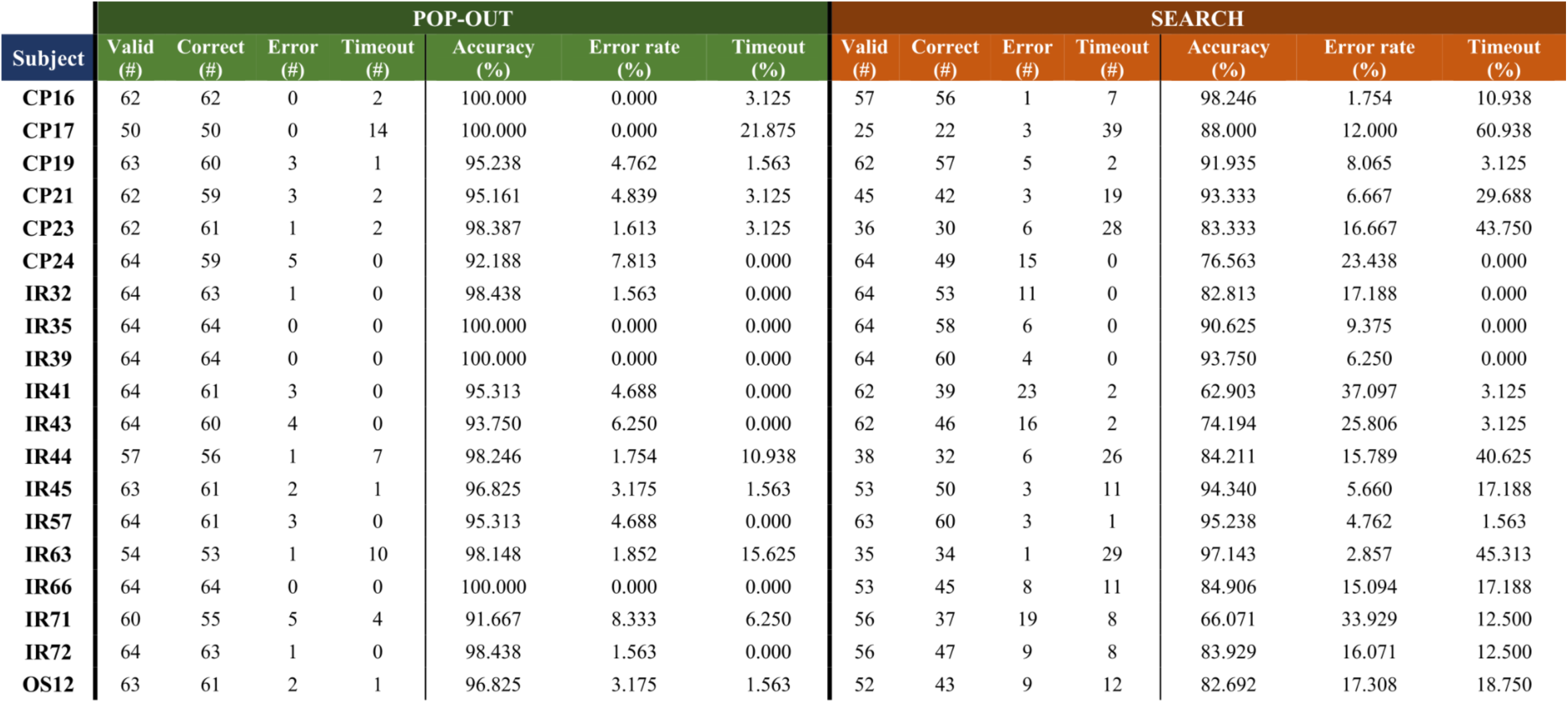
Individual participant behavioral performance by condition. The table reports for each participant the number (*#*) of valid, correct, error, and time-out trials, together with the accuracy (% correct over responded trials), error rate (1 – accuracy), the percentage of time-out trials.

### SI – 2. Power analyses

#### SI – 2.1 Details for the nonparametric cluster-based permutation test

Time-frequency power estimates in each condition were estimated using a Morlet wavelet transform. Between-condition differences were assessed using a nonparametric cluster-based permutation approach over frequencies 1–150 Hz and time points in post-stimulus interval [0, +2.0] s (see results in Fig. 1D). Details about the regions contributing to the observed differences are provided in the Supplementary Table 2 (shown below). Twelve ROIs showed significant clusters in both directions.

**Supplementary Table 2.** Regions of interest contributing significant clusters in the Search vs. Popout power contrast (cluster-based permutation test, *p_perm_*<0.05).

| Hemisphere | Higher power in Search | Higher power in Pop-out |
| --- | --- | --- |
| Left | IFS, **MFG**, **MTG**, **MTL**, **OFC**, **OG**, PMC, SFG, SMA, STG, STS, TP, medial PFC | Amy, FFG, IFS, IPL, ITG, **MFG**, **MTG**, **MTL**, MTS, **OFC**, **OG**, **PMC**, **SMA**, **STG** |
| Right | ACC, **MCC**, **MFG**, **MTL**, OFC, OG | IFG, IPL, M1, **MCC**, **MFG**, **MTG**, **MTL**, PMC, SMA, SPL, TPJ |

*Highlighted in bold are regions showing significant differences in both directions: Left IFS, MFG, MTG, MTL, OFC, OG, PMC, SMA, STG; Right MCC, MFG, MTL.*

#### SI – 2.2 Site-resolved test for early HFA effects

The main-text HFA analyses characterize condition differences at the level of anatomically pooled regions of interest (ROIs). To test whether this pooling could obscure early, pop-out-specific effects present at only a subset of ventral recording sites, as previously reported in fusiform cortex (Ossandón et al., 2012), we performed a complementary analysis in which the single contact, rather than the ROI, was the unit of analysis. The analysis was restricted a priori to ventral occipito-temporal cortex, the region in which Ossandón et al. reported early pop-out effects (their condition-differing sites were predominantly temporal and occipital, and fusiform-heavy; only 11 of 123 transient sites were parietal). We therefore examined all contacts assigned to occipital gyrus (OG), fusiform gyrus (FFG), and inferior temporal gyrus (ITG). This yielded 109 ventral contacts (OG, 43; FFG, 29; ITG, 37) across 9 of the 19 participants entering the HFA analyses. Parietal cortex was not included because the prior effect was localized to ventral occipito-temporal sites.

For each contact we used single-trial HFA (70–150 Hz), computed and baseline-normalized as in the main analysis (z-scored relative to a pre-stimulus baseline of -2.0 to -1.5 s). Before comparing conditions, each contact had to pass a responsiveness gate designed to ensure that a Pop-out versus Search difference was evaluated only where a genuine task-evoked response was present: the contact had to show at least one significant positive HFA cluster relative to baseline over the 0–1000 ms post-stimulus window in at least one condition (cluster-based permutation test, cluster-defining threshold *p*<0.05, 10,000 permutations), with no minimum-duration requirement so that early and transient responders of the kind described by Ossandón et al. were retained. Of the 109 ventral contacts, 39 passed this gate (OG, 17/43; FFG, 13/29; ITG, 9/37) and entered the condition comparison.

For each responsive contact, HFA was compared between Pop-out and Search using a cluster-based permutation test across single trials (independent-samples *t*-statistic at each time sample; cluster-defining threshold *p*<0.05, two-tailed; cluster mass as the cluster-level statistic; 10,000 permutations of condition labels; fixed random seed). To restrict the test to the pre-response processing phase in both conditions and thereby avoid a response-bounded duration confound, the comparison window was set to *0* to *T_end*, with *T_end* = min(group-mean Pop-out reaction time, 500 ms). Because the group-mean Pop-out reaction time was 774 ms, the window was selected as 0–500 ms for all contacts. Surviving clusters were recorded with their direction (Pop-out > Search or Search > Pop-out), onset latency, peak latency, cluster mass, and cluster-level *p*-value. A significant cluster was classified as early if its onset preceded 300 ms.

As a like-for-like replication of the Ossandón et al. contrast, we additionally computed, per contact, the mean HFA over their 100–500 ms window on each trial and compared conditions with a Wilcoxon rank-sum test, correcting across responsive contacts with the Benjamini-Hochberg false-discovery-rate procedure. Finally, we evaluated the site-level results against chance at the group level. For each ROI and for the pooled ventral sample, the observed number of significant contacts was compared to the number expected under the 5% false-positive rate (0.05 × number of responsive contacts) using a binomial test, and the directional split among significant contacts was compared to an even split using a binomial test against 0.5.

Of the 109 ventral contacts, 39 (35.8%) passed the responsiveness gate and entered the Pop-out versus Search comparison (OG, 17/43; FFG, 13/29; ITG, 9/37). Among these 39 responsive contacts, 7 showed a significant Pop-out versus Search difference in HFA, more than the ∼1.95 expected under the 5% false-positive rate (binomial test, *p*=0.0029), indicating genuine site-level heterogeneity that is resolvable at the single-contact level but averaged out in the pooled ROI analysis. These effects were concentrated in the fusiform cortex (Supplementary Fig. 1A,B). Significant contacts exceeded chance in FFG (4/13 responsive; binomial *p*=0.0031) but not in OG (2/17) or ITG (1/9), neither of which departed from chance individually. Their direction was predominantly Pop-out > Search (5 of 7 contacts). The confirmatory analysis over the 100–500 ms window (per-trial mean HFA, FDR-corrected across responsive contacts) yielded a consistent picture: 7 contacts were significant, 6 in the Pop-out > Search direction and predominantly fusiform (left OG IR45/TPG4, z=6.59; left FFG IR45/TPG34, z=5.19; left FFG IR45/TPG26, z=4.52; left ITG IR72/LBT_6_7, z=3.30; left FFG IR72/LBT_5_6, z=3.14; left FFG IR72/RTH_4_5, z=2.98), with a single occipital site in the opposite direction (left OG IR45/TPG18, z=-2.68). A subset of these fusiform effects had early onsets: three of the seven significant contacts showed a cluster onset before 300 ms, and two before 200 ms. The clearest example was a left fusiform site with an early (∼200 ms), transient response that was stronger in Pop-out than in Search (IR45/TPG26; *p_cluster_*=0.0002; Supplementary Fig. 1C, left), accompanied by occipital and fusiform sites with stronger and more sustained Pop-out > Search responses (IR45/TPG4 and IR45/TPG34; Supplementary Fig. 1C, center and right). A second fusiform site showed the same early direction with a slightly later onset (CP17/RMT3; ∼252 ms), while one early site showed the opposite sign (left FFG CP23/LMTP2; Search > Pop-out, ∼120 ms), indicating that the ventral early effects are heterogeneous in sign.

Overall, these significant contacts were a small minority of the responsive ventral sample (7/39) and of all ventral contacts (7/109), arising from 4 of the 9 participants. Their spatial restriction to a handful of fusiform and occipital sites, together with their transient time course, is consistent with sparse, localized bottom-up signals at select ventral sites rather than a spatially extended process engaged uniformly across conditions.

**Supplementary Fig. 1.**
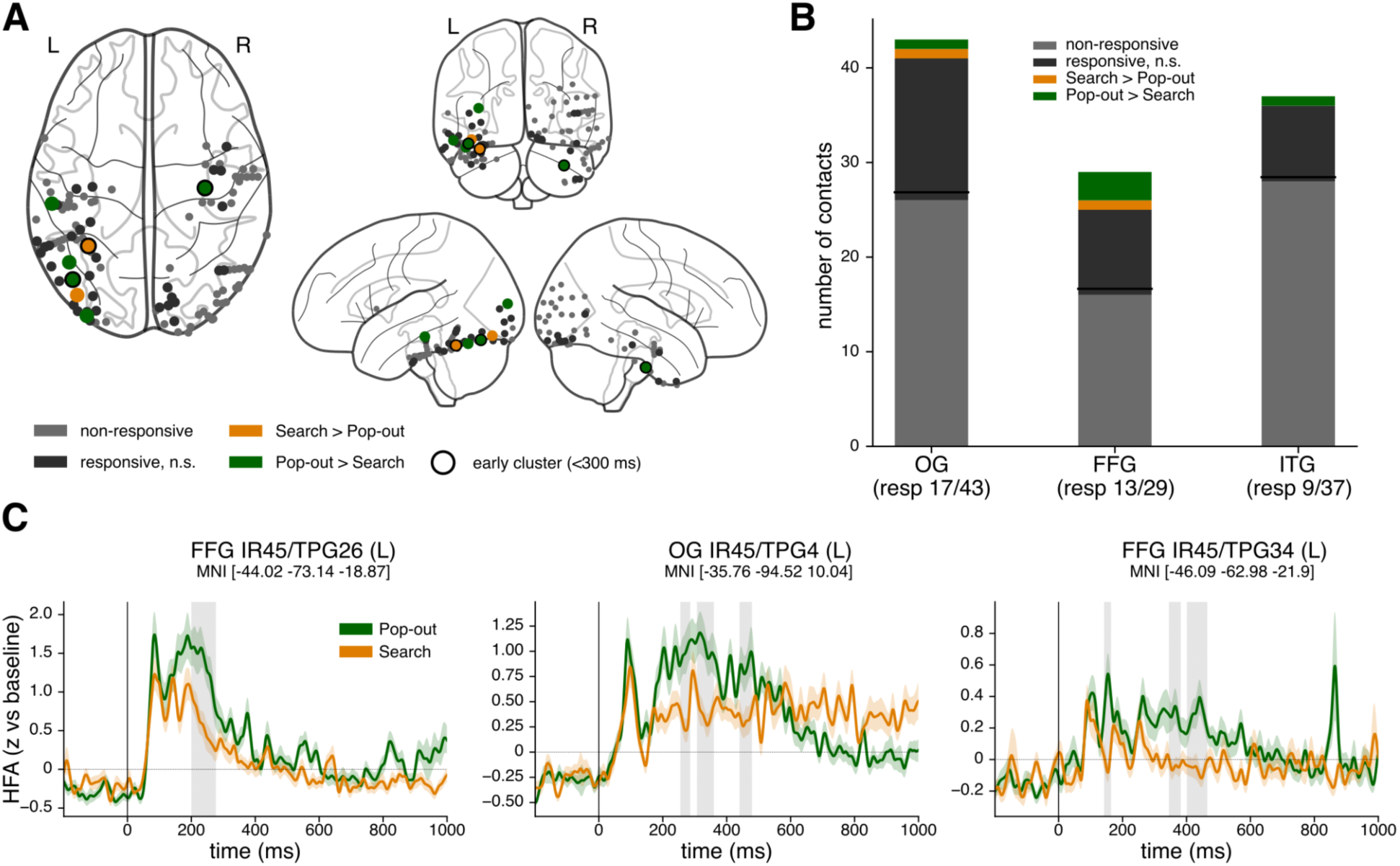
Site-resolved test for early HFA effects in ventral occipito-temporal cortex. **A**, All ventral contacts displayed on a template glass brain (axial, coronal, and left/right sagittal views; L, left; R, right). Contacts are colored by outcome: non-responsive (light grey) and responsive but non-significant (dark grey) sites did not show a significant Pop-out vs Search difference; responsive sites with a significant difference are colored by direction (Pop-out > Search, green; Search > Pop-out, orange). Black rings mark contacts whose significant cluster had an early onset (<300 ms). **B**, Contact counts per ROI (OG, FFG, ITG), stacked into non-responsive, responsive non-significant, and significant-by-direction (Search > Pop-out, orange; Pop-out > Search, green); the number of responsive contacts is given below each bar. **C**, Three representative responsive contacts (mean ± SEM across trials; Pop-out, green; Search, orange; z-scored to baseline; t = 0, search-display onset). Grey shading marks time windows of significant Pop-out vs Search difference (cluster-based permutation, *p*<0.05). Left: a fusiform site (FFG, IR45/TPG26) with an early (∼200 ms), transient Pop-out > Search response of the kind reported by Ossandón et al. (2012). Center: an occipital site (OG, IR45/TPG4) with a stronger and more sustained Pop-out > Search response. Right: a fusiform site (FFG, IR45/TPG34) with a later, sustained Pop-out > Search difference. MNI coordinates are given above each panel.

### SI – 3. Dynamic time warping (DTW)

#### SI – 3.1 Time warping using mismatched ROI pairs as null

Because dynamic time warping (DTW) is designed to find the optimal temporal alignment between any two signals, a significantly positive warping offset during the task interval demonstrates that Search and Pop-out can be temporally remapped onto one another, but does not by itself establish that they share a common underlying trajectory. To test for a genuine shared trajectory, we asked whether the empirical, same-region Search–Pop-out pairing becomes more similar, and aligns at lower cost, than DTW can achieve for unrelated signals. For each of the eleven ROIs that passed both validity criteria (’valid’ group), we computed a matched alignment between the empirical Search and Pop-out high-frequency activity (HFA) of that ROI, and a mismatched null in which the Search trace of the valid ROI was warped onto Pop-out traces drawn from ROIs that did not pass the validity criteria (’invalid’ group; e.g., Left OG Search vs. Left SFG Pop-out). Over the post-stimulus task interval we quantified four metrics: pre-warp similarity (Pearson correlation between Search and Pop-out before warping), post-warp similarity (correlation after DTW alignment), similarity change (post-minus pre-warp similarity), and alignment cost (the integral warping offset, indexing the amount of temporal remapping required). Matched and mismatched distributions were compared per ROI (Mann–Whitney *U* test, *p*<0.05; Supplementary Fig. 2 and Supplementary Table 3).

In every ROI, Matched pairs showed significantly higher pre- and post-warp similarity and significantly lower alignment cost than Mismatched pairs (all *p*<0.05; exact values in Supplementary Table 3). Before warping, matched pairs were already more correlated than mismatched ones (matched pre-warp similarity 0.11–0.64 across ROIs, versus -0.16 to 0.09 for mismatched; all *p*<0.05, largest *p*=0.0350 in Right ACC), indicating that same-region Search and Pop-out signals share temporal structure independently of any warping. After warping, matched pairs reached substantially higher similarity than mismatched pairs (post-warp similarity 0.91–0.97 versus 0.73–0.87; all *p*<0.05, largest *p*=0.0112 in Right TPJ), and required less temporal distortion to do so (alignment cost lower for matched in every ROI; all *p*<0.05, largest *p*=0.0208 in Left OG). Thus, in each of the eleven ROIs, the empirical Search and Pop-out traces are both more alike to begin with and more closely aligned after optimal warping, at lower cost, than DTW produces for unrelated signals.

Similarity change was by contrast larger for mismatched than the matched pairs in ten of the eleven ROIs (mismatched change 0.75–1.01 versus matched 0.33–0.80; the exception was Right ACC, where the two were comparable). This pattern is expected and is informative rather than contradictory. Because matched pairs already begin with high similarity, there is comparatively little room for warping to improve their alignment; whereas mismatched pairs begin near zero correlation and can therefore be improved more (e.g., Left OG: +0.87 for mismatched versus +0.33 for matched). Critically, this confirms that DTW is indeed able to substantially increase the apparent similarity of unrelated signals, which is the very flexibility that motivates this control. Yet even after this larger correction, and despite the greater amount of warping applied, the mismatched pairs do not approach the post-warp similarity, nor the low alignment cost, of the matched pairs. The greater improvability of mismatched signals therefore does not undermine but underscores the specificity of the matched result: warping cannot manufacture, out of unrelated regions, the degree of correspondence observed for genuine same-region Search–Pop-out pairs.

Together, these analyses demonstrate that the temporal alignment between Search and Pop-out reflects a genuine shared neural trajectory rather than the trivial alignability of two task-evoked signals.

**Supplementary Table 3.**
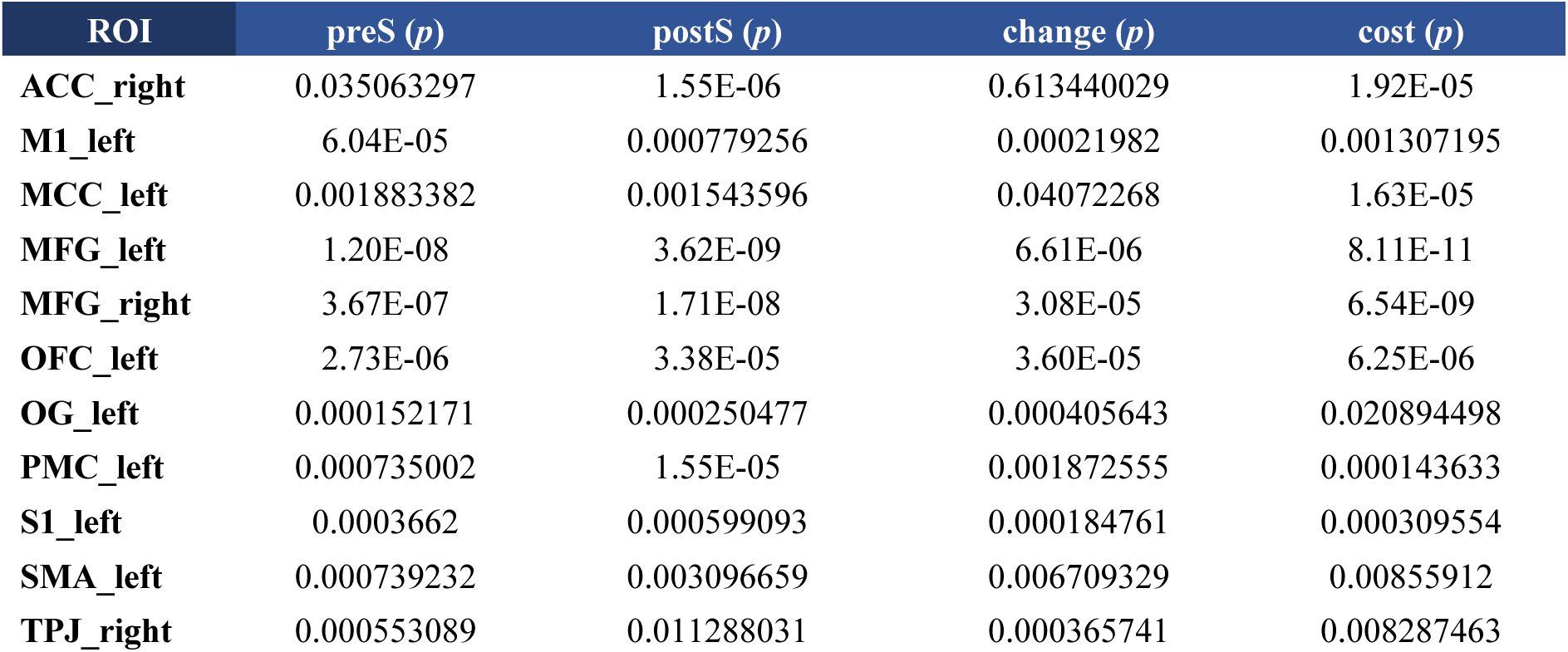
Statistical comparison of matched versus mismatched DTW alignment across the eleven valid ROIs. For each ROI that passed both DTW validity criteria, *p*-values are reported for the matched-versus-mismatched comparison on four metrics computed over the post-stimulus interval: *pre-warp similarity* (’preS’), *post-warp similarity* (’postS’), *similarity change* (’change’), and *alignment cost* (’cost’).

| ROI | preS (*p*) | postS (*p*) | change (*p*) | cost (*p*) |
| --- | --- | --- | --- | --- |
| ACC_right | 0.035063297 | 1.55E-06 | 0.613440029 | 1.92E-05 |
| M1_left | 6.04E-05 | 0.000779256 | 0.00021982 | 0.001307195 |
| MCC_left | 0.001883382 | 0.001543596 | 0.04072268 | 1.63E-05 |
| MFG_left | 1.20E-08 | 3.62E-09 | 6.61E-06 | 8.11E-11 |
| MFG_right | 3.67E-07 | 1.71E-08 | 3.08E-05 | 6.54E-09 |
| OFC_left | 2.73E-06 | 3.38E-05 | 3.60E-05 | 6.25E-06 |
| OG_left | 0.000152171 | 0.000250477 | 0.000405643 | 0.020894498 |
| PMC_left | 0.000735002 | 1.55E-05 | 0.001872555 | 0.000143633 |
| S1_left | 0.0003662 | 0.000599093 | 0.000184761 | 0.000309554 |
| SMA_left | 0.000739232 | 0.003096659 | 0.006709329 | 0.00855912 |
| TPJ_right | 0.000553089 | 0.011288031 | 0.000365741 | 0.008287463 |

**Supplementary Fig. 2.**
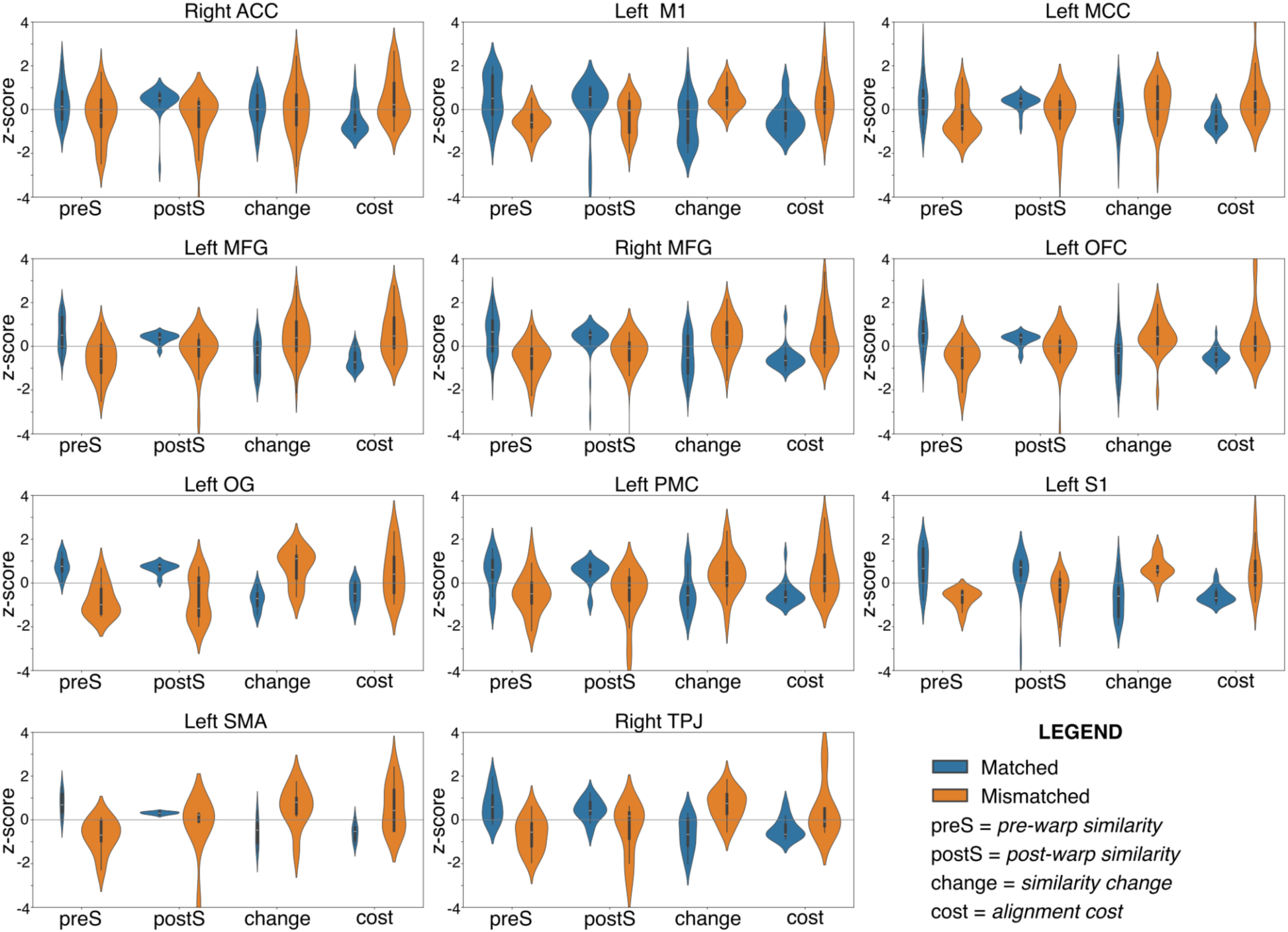
A mismatched-ROI null confirms a shared Search–Pop-out neural trajectory in the eleven valid ROIs. For each of the ROIs that passed both DTW validity criteria (one panel per ROI), we compared the empirical, same-region pairing of Search and Pop-out high-frequency activity (’Matched’, blue) against a null in which the Search trace of the valid ROI was warped onto Pop-out traces drawn from ROIs that did not pass the validity criteria (’Mismatched’, orange). Violin plots show the distribution, across all recording sites contributing to each ROI, of four metrics computed over the post-stimulus task interval: *pre-warp similarity* (’preS’), *post-warp similarity* (’postS’), *similarity change* (’change’), and *alignment cost* (’cost’). All metrics are z-scored across the pooled Matched and Mismatched values within each ROI for display on a common axis. Within each violin plot, the black box plot represents 95%, 75%, 50%, 25%, and 5% percentile.

#### SI – 3.2 Quantification of warping path geometry

Across all eleven ROIs that passed both validity criteria, the task-segment warping function was well approximated by a straight line (mean linear R² = 0.920, range 0.857–0.955), indicating that the temporal scaling between Pop-out and Search is predominantly uniform (Supplementary Table 4). The instantaneous local slope showed only a weak, uniformly positive trend across the task segment (slope-vs-time correlation 0.010–0.073 across ROIs), and the warping-offset humps were close to symmetric (skewness between -0.291 and +0.130, with inconsistent sign across regions) (Supplementary Fig. 3). Together these measures indicate that the offset humps arise principally from approximately uniform stretching, with at most a mild and spatially inconsistent tendency toward greater stretching later in the task segment. Left OG, which appeared atypically late-concentrated in the single-region example (Fig. 2D), had the lowest linear R² of the valid regions but was nonetheless well fit by a straight line, and its offset hump was essentially symmetric (skewness = –0.028) rather than late-skewed; its apparent non-uniformity therefore does not survive quantification. A mid-segment offset peak, which DTW produces under either uniform or stage-specific scaling, is not by itself diagnostic. The near-linear warping functions and near-symmetric humps are what adjudicate in favor of uniform dilation.

**Supplementary Table 4.**
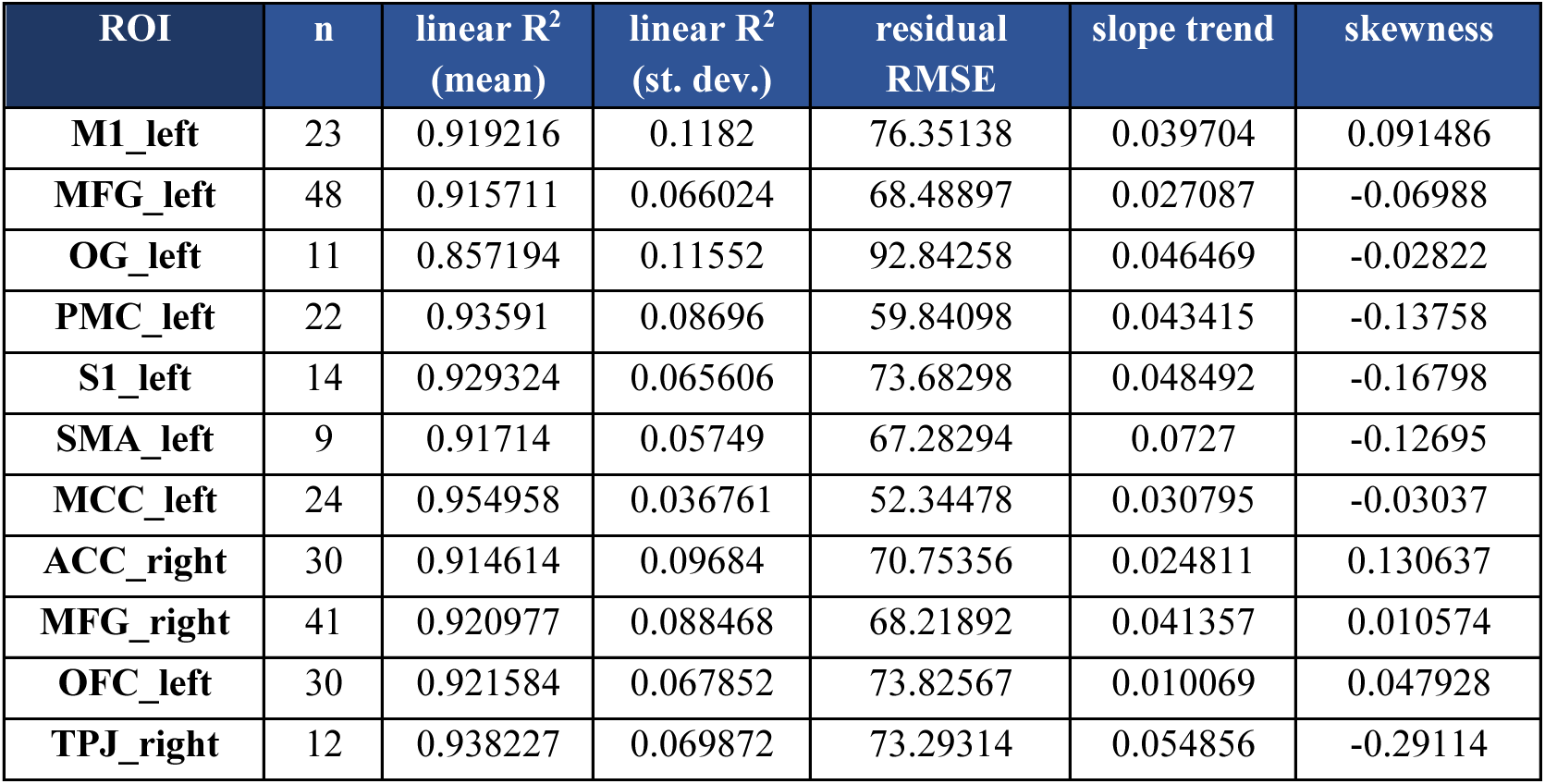
Quantification of warping-path geometry. For each ROI that passed both DTW validity criteria, the table reports the number of electrode contacts (*n*) together with the following geometry metrics: the linear R² coefficient (mean and standard deviation); residual RMSE of a straight-line fit; the slope trend; and the skewness of the warping offset.

**Supplementary Fig. 3.**
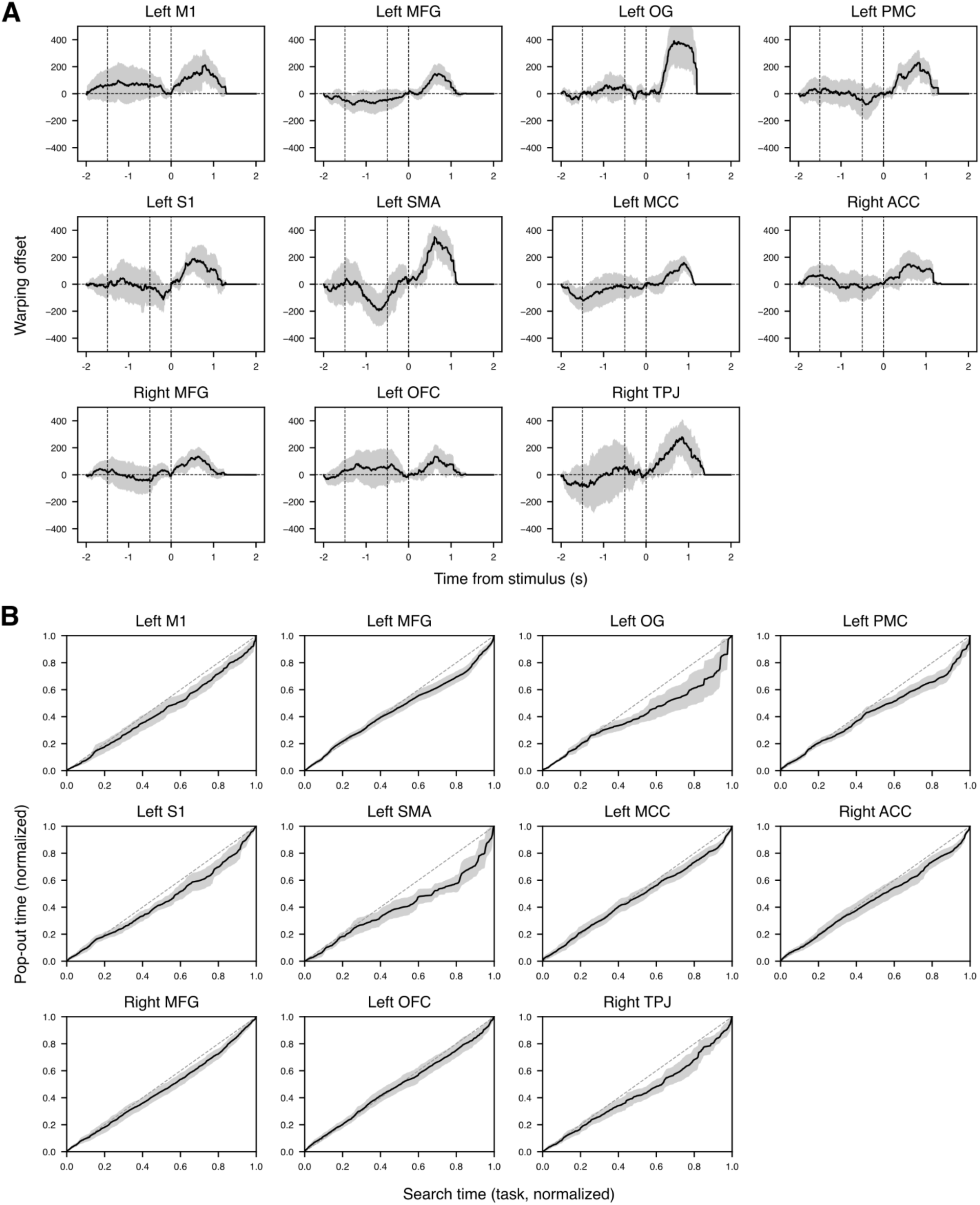
Geometry quantification reveals that temporal scaling is predominantly uniform. **A**, Warping offset curve (Pop-out - Search) for each of the eleven valid ROIs. **B**, Warping function. Shaded band indicates 95% confidence interval.

#### SI – 3.3 Interval-matched control

Two control analyses were performed to test whether the temporal-scaling relationship between conditions reflects a general dependence of the neural trajectory on response speed: i) Reaction-time (RT)-matched control, ii) Within-condition Fast vs. Slow.

**i) RT-matched control.** To test whether the temporal-scaling relationship between Pop-out and Search survives once trial duration is controlled, we repeated the DTW analysis on RT-matched subsets of trials. Within each participant we constructed RT-matched Pop-out and Search subsets by histogram stratification: the two conditions’ RTs were pooled and partitioned into fixed-width bins (50 ms, anchored at multiples of the bin width across the pooled range), and within each bin we retained min(n_Pop-out, n_Search) trials per condition by random subsampling without replacement (fixed seed). Bins containing trials from only one condition contributed nothing, which removed the non-overlapping fast-Pop-out and slow-Search tails. Matching was performed strictly within participants, with pooling across participants only afterwards. Participants retaining fewer than 16 trials per condition were excluded. Match quality was assessed per participant with a two-sample Kolmogorov–Smirnov test, the standardized mean difference (SMD), and the overlap coefficient of the two RT densities; a participant was retained only if the matched RT distributions were statistically indistinguishable (KS *p*>0.05) and equivalent (|SMD|<0.1). For each retained participant, single-trial HFA was normalized by a per-trial baseline z-score and averaged over the matched trials to form the per-electrode series, for all electrodes. Because matching equates the two conditions’ RT distributions, both matched series were truncated at the participant’s mean RT computed over the pooled matched trials (matched Pop-out and matched Search combined), applied identically to both series. The matched series were then passed unchanged through our DTW pipeline. ROI validity was assessed on the matched results exactly as in the original pipeline, requiring both the absence of a significant pre-stimulus offset and a significantly positive task-segment warping offset (see Methods). We further quantified alignment on the task (post-stimulus) segment with two metrics of the warping offset (task-segment peak and integral offset). Each empirical metric was compared, per ROI and pooled, against a null distribution generated by randomly relabeling the matched trials as Pop-out or Search within each participant (preserving per-condition subset sizes), re-averaging, and re-running the identical DTW pipeline and truncation (1,000 permutations); we report each empirical value’s percentile and a two-sided permutation *p*-value.

Within-participant histogram stratification produced RT-matched subsets whose distributions were statistically indistinguishable between conditions (all retained participants KS *p*>0.98, |SMD|<0.06, overlap 0.71–0.90) (Supplementary Fig. 4A). The binding constraint was trial count rather than match quality, with one participant excluded for retaining fewer than 16 trials per condition. After matching, the average HFA amplitude profiles of Pop-out and Search overlapped closely in every ROI (Supplementary Fig. 5), indicating that RT-matching removed the gross temporal differences between conditions.

Correspondingly, the DTW warping offset collapsed toward zero relative to the unmatched analysis (Supplementary Fig. 4B). We re-assessed ROI validity on the RT-matched results using the same two criteria as the original pipeline. After RT-matching, none of the eleven regions that were valid in the original analysis retained a significantly positive warping offset during the task segment (Supplementary Table 5): the one-sided task-segment offset *t*-test was non-significant in every ROI (all *p*>0.15), so no ROI (0/11) met the joint validity criteria. This was specifically a loss of the positive task-segment offset (i.e., the signature of temporal scaling), rather than a pre-stimulus artifact: the pre-stimulus variance criterion was still satisfied in ten of eleven regions (Left OFC being the only exception). By contrast, the same regions were all valid in the original, duration-uncontrolled analyses. Thus the very feature that defined these ROIs as showing temporal scaling between Pop-out and Search disappears once trial duration is equated.

Across the two warping offset metrics, the empirical values fell within the within-participant condition-label-shuffle null for the pooled analysis and for every ROI (Supplementary Fig. 4C): the pooled task-segment peak offset (*p*=0.820) and integral (*p*=0.490) were indistinguishable from chance relabeling, and no ROI’s offset exceeded the null at *p*<0.05 (Supplementary Table 6). Two regions showed peak offsets trending above the null, but without reaching significance (Left OG and Left S1, *p*=0.062 for the offset peak only, not the offset integral). These results indicate that the Pop-out↔Search temporal-scaling effect observed in the original analysis is accounted for by the difference in trial duration: when RT is held constant by matching, no residual condition-specific trajectory difference survives.

**Supplementary Table 5.** DTW analysis on reaction-time (RT)-matched Pop-out and Search. Results of the DTW analysis between RT-matched conditions, reporting the two validity criteria for pre-stimulus offset (*Seg1 p*) and warping offset during the task segment (*Seg2 p*), together with the measures of peak warping offset, time to peak, and integral offset.

| ROI | Seg1 p | Seg2 p | Peak | Time (to peak) | Integral |
| --- | --- | --- | --- | --- | --- |
| ACC_right | 0.655994 | 0.303916 | 47.86667 | 894 | 8017.817 |
| M1_left | 0.712506 | 0.435389 | 36.56522 | 499 | -678.087 |
| MCC_left | 0.808859 | 0.912154 | 10.83333 | 14 | 86.77778 |
| MFG_left | 0.083962 | 0.870752 | 16.6125 | 12 | 174.9625 |
| MFG_right | 0.134681 | 0.667175 | 15.84286 | 976 | -10856 |
| OFC_left | 0.016424 | 0.848107 | 8.703704 | 382 | -1992.33 |
| OG_left | 0.448144 | 0.221365 | 133.5909 | 701 | 883.2727 |
| PMC_left | 0.980949 | 0.154912 | 59.52273 | 766 | 20009.23 |
| S1_left | 0.976443 | 0.170464 | 112.3571 | 756 | 28732.61 |
| SMA_left | 0.06848 | 0.824262 | 25.5 | 918 | -45928.3 |
| TPJ_right | 0.099749 | 0.384631 | 59.625 | 822 | 6791.583 |

*Highlighted in orange background is for each ROI the criterion that did not pass statistical threshold.*

**Supplementary Table 6.** Statistical comparison of RT-matched Pop-out versus Search DTW alignment across the eleven valid ROIs. Results of the control analysis comparing each empirical metric (peak/integral offset) against a null distribution, obtained from a within-participant condition-label-shuffle (1,000 permutations; two-sided *p*).

| ROI | Peak (empirical) | Peak (null mean) | Peak (percentile) | Peak (p) | Integral (empirical) | Integral (null mean) | Integral (percentile) | Integral (p) |
| --- | --- | --- | --- | --- | --- | --- | --- | --- |
| M1_left | 36.56522 | 40.85513 | 48.5 | 0.864 | -678.087 | -234.219 | 33 | 0.927 |
| MFG_left | 16.6125 | 31.94209 | 27.5 | 0.509 | 174.9625 | -1201.83 | 41.8 | 0.852 |
| OG_left | 133.5909 | 60.78091 | 93.9 | 0.062 | 883.2727 | 1224.898 | 44.1 | 0.948 |
| PMC_left | 59.52273 | 45.6208 | 70.8 | 0.689 | 20009.23 | 388.9217 | 87.1 | 0.282 |
| S1_left | 112.3571 | 53.44932 | 93.9 | 0.062 | 28743.61 | -928.863 | 95.1 | 0.155 |
| SMA_left | 25.5 | 64.90244 | 20.8 | 0.394 | -46014.8 | -1146.67 | 7 | 0.092 |
| MCC_left | 10.83333 | 44.71214 | 12.3 | 0.277 | 86.77778 | -328.679 | 40.8 | 0.913 |
| ACC_right | 48.73333 | 37.00365 | 72.5 | 0.629 | 8007.217 | -625.819 | 77 | 0.434 |
| MFG_right | 15.84286 | 36.5602 | 23.8 | 0.439 | -10476.1 | -1188.23 | 19.1 | 0.413 |
| OFC_left | 8.703704 | 42.69631 | 7.9 | 0.198 | -1992.33 | -1867.2 | 32.9 | 0.996 |
| TPJ_right | 59.625 | 66.45363 | 48.8 | 0.895 | 6791.417 | -209.933 | 58.4 | 0.655 |
| POOLED | 11.42708 | 13.83883 | 47.3 | 0.820 | -3040.87 | -323.509 | 21.6 | 0.490 |

*Results that did not exceed the null at $p < 0.05$ are highlighted in orange background for each ROI.*

**Supplementary Fig. 4.**
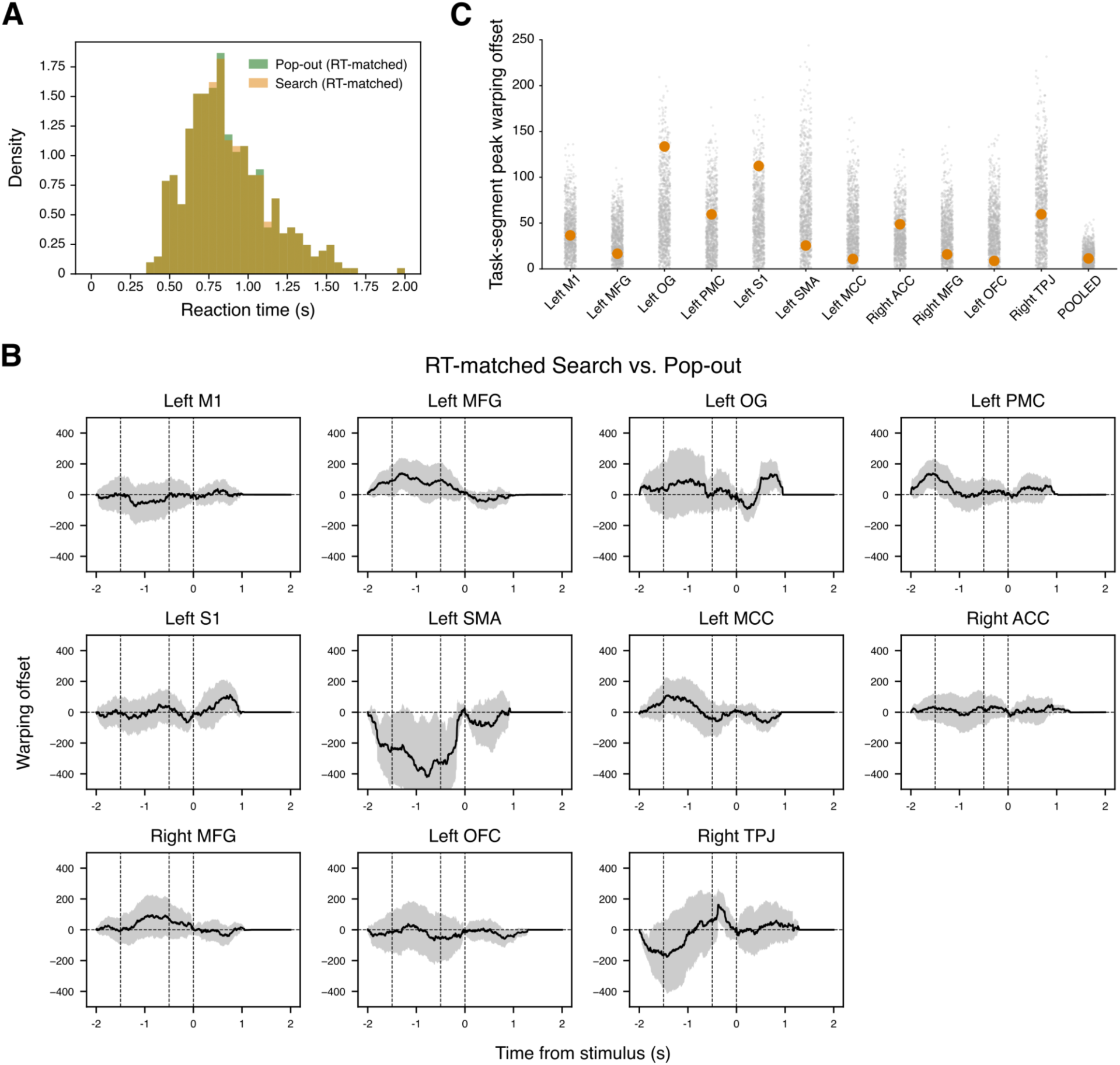
RT-matched control reveals that the temporal scaling effect disappears when equating trial duration between conditions. **A**, Reaction time (RT) distributions pooled across all participants for Pop-out (green) and Search (orange), after stratification. **B**, Warping offset curve (RT-matched Pop-out - Search) for each of the eleven valid ROIs. **C**, Task-segment peak warping offset is shown for each valid ROI comparing the empirical value (orange) against a null distribution (each gray dot represents one permutation).

**Supplementary Fig. 5.**
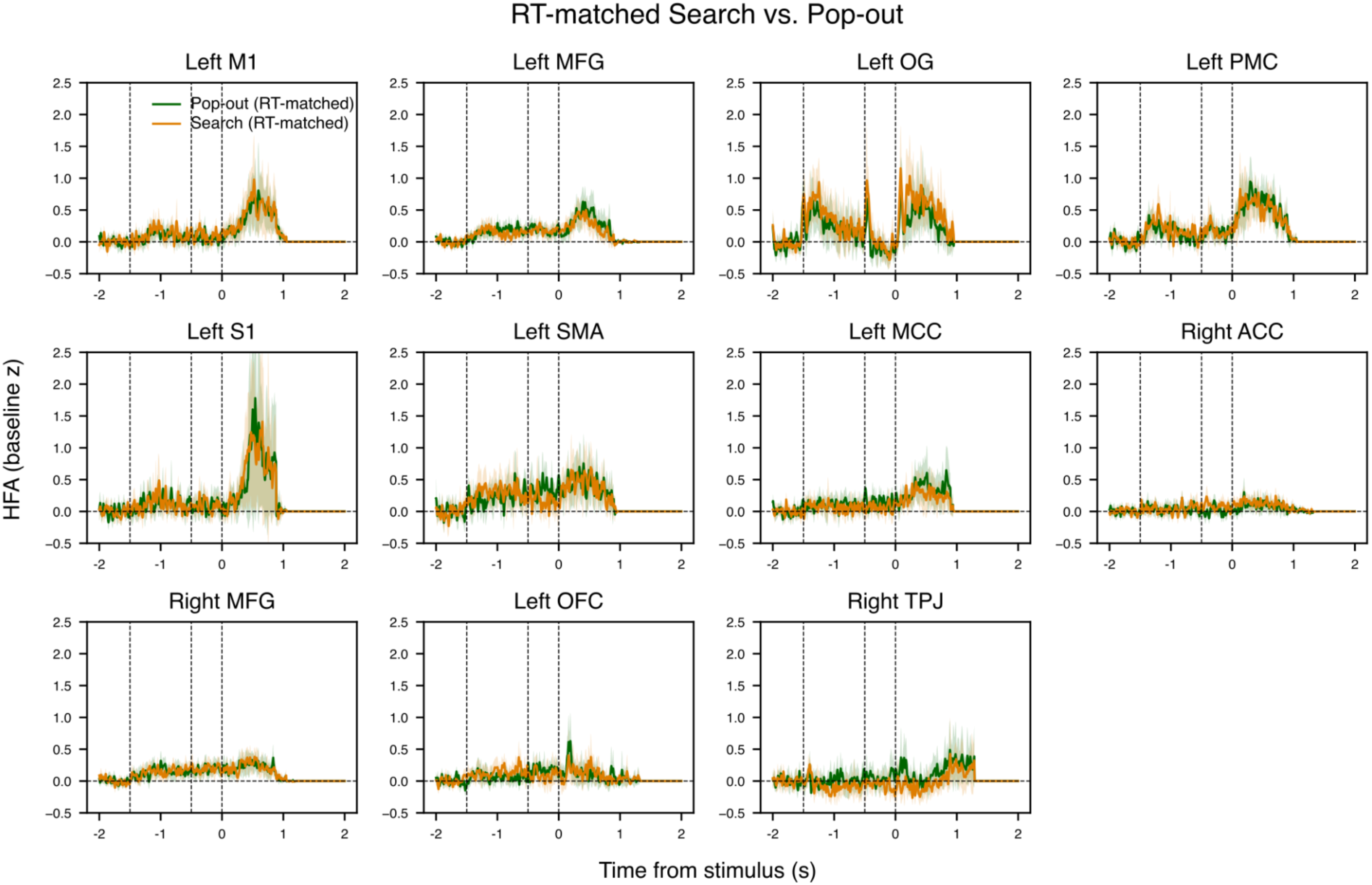
RT-matched conditions show overlapping HFA amplitude profiles in every valid ROI. Trial-averaged HFA signals (z-scored), shown for Search (orange) and Pop-out (green), are closely overlapping after RTs stratification between conditions.

#### SI – 3.4 Within-condition Fast vs. Slow

**ii) Within-condition Fast vs. Slow.** To test whether the temporal-scaling relationship between conditions reflects a general dependence of the neural trajectory on response speed, we asked whether the same scaling arises within a single condition between faster and slower trials. Within each condition (Search; Pop-out) and each participant, trials were divided into Fast and Slow subsets by a median split on single-trial reaction time (RT): trials were rank-ordered by RT and the lower and upper halves were assigned to Fast and Slow, with the single middle trial dropped when the trial count was odd so that the two subsets had equal size. Participants retaining fewer than 16 trials per subset were to be excluded; all participants met this floor in both conditions. For each participant, single-trial HFA was normalized by a per-trial baseline z-score and averaged over the Fast and Slow subsets separately to yield the per-electrode series, for all electrodes. Because Fast and Slow subsets differ in duration by design, both averaged series were truncated at the participant’s mean RT of the Slow subset, computed separately within each condition, mirroring the original pipeline’s use of the slower condition’s mean RT; in the permutation null this truncation point was held fixed at each participant’s real Slow-subset mean. The Fast and Slow series were passed unchanged through our DTW pipeline, with the Slow series treated as the reference so that a positive task-segment warping offset indicates that Slow is a temporally stretched version of Fast. Before warping, we assessed early-window differences by comparing the mean HFA amplitude over 0–300 ms post-stimulus between Fast and Slow with a two-sided paired *t*-test across contacts, per ROI. ROI validity was evaluated with the original two criteria (one-sided χ² pre-stimulus variance test, null variance ≤ 300²; one-sided one-sample *t*-test of a positive task-segment offset). On the task segment we quantified the Fast-vs-Slow alignment with two metrics of the warping offset (task-segment peak and integral). We compared each metric, per ROI and pooled, against a null generated by randomly relabeling each participant’s retained trials into two pseudo-groups of the same sizes as the real Fast/Slow subsets, re-averaging, and re-running the identical pipeline and truncation (1,000 permutations). Because the hypothesis is that a genuine within-condition speed difference produces more alignment than chance relabeling, we report the one-sided upper-tail permutation *p*-value (the probability that the null equals or exceeds the empirical value) together with the empirical percentile. Search and Pop-out were analyzed independently.

Within each condition, the median split produced Fast and Slow subsets of comparable size (22– 32 trials each) for all participants. The Fast and Slow HFA trajectories did not differ in the early post-stimulus window (the 0–300 ms amplitude comparison was non-significant in any ROI for either Search or Pop-out), indicating that the two subsets were matched on early activation and differed principally in the temporal extent of the response.

Dynamic time warping revealed a positive task-segment warping offset between Fast and Slow trials within condition, with five of the eleven ROIs that were valid in the original analysis meeting both validity criteria in each condition (Supplementary Table 7). Within Search, 5 ROIs retained a significantly positive warping offset during the task segment (Left M1, Left MFG, Left SMA, Left OG, and Right MFG), and 2 ROIs showed offsets trending above the null, but without reaching significance (Left PMC and Left OFC, Segment 2: *p*=0.0626 and *p*=0.0692, respectively). Within Pop-out, 5 ROIs retained a significantly positive warping offset during the task segment (Left M1, Left MFG, Left SMA, Left PMC, Left MCC), in addition to Left OG showing warping offset trending to significance but not reaching it (Left OG, Segment 2: *p*=0.0515). Further, the warping offset metrics exceeded the within-condition Fast/Slow-label-shuffle null: the pooled task-segment peak offset and integral both exceeded the null (*p*<0.001) in Search and in Pop-out (Supplementary Table 8). At the regional level, the peak warping offset exceeded the null in 7 of 11 ROIs in Search and 8 of 11 in Pop-out, with 6 ROIs robust in both conditions (Left M1, Left MFG, Left OG, Left PMC, Left SMA, and Right TPJ) (Supplementary Fig. 6 and Supplementary Table 8). For the integral offset, 5 of 11 ROIs exceeded the null in Search and 6 in Pop-out (robustly in Left M1, Left MFG, Left PMC, and Left SMA) (Supplementary Table 8).

We note that, in both conditions, the evidence of temporal scaling at the single-ROI level is weaker in this within-condition analysis compared to the original analysis. This may be due to lower statistical power in some ROIs, attributable to a reduction in the number of trials for each compared subset (Fast, Slow), after splitting trials in two halves within each task condition (Search, Pop-out) based on RTs, which likely accounts for the less consistent significance of the alignment cost. The pooled results, however, are unambiguous in both conditions, and across offset metrics. Further, the ROIs that robustly showed a temporal scaling effect belong to the group of areas showing a robust task-evoked HFA response (i.e., Left M1, MFG, OG, PMC, and SMA). These results indicate that a within-condition temporal scaling is present in both Search and Pop-out: slower trials trace the same neural trajectory as faster trials, stretched in time. Together with the between-condition scaling and its dependence on trial duration, this supports the interpretation that a single cortical mechanism generates the visual-search response.

**Supplementary Table 7.** DTW analysis on within-condition Fast vs. Slow. Results of the DTW analysis between Fast and Slow trials in each condition, separately, reporting the two validity criteria for pre-stimulus offset (*Seg1 p*) and warping offset during the task segment (*Seg2 p*), together with the measures of peak warping offset, time to peak, and integral offset.

|  | ROI | Seg1 p | Seg2 p | Peak | Time (max) | Integral |
| --- | --- | --- | --- | --- | --- | --- |
| SEARCH | ACC_right | 0.18542235 | 0.39718411 | 58 | 1232 | 3625.43333 |
|  | M1_left | 0.41853381 | 0.00280336 | 237.978261 | 948 | 68494.3043 |
|  | MCC_left | 0.0192182 | 0.10186333 | 96.6875 | 928 | 44358.4583 |
|  | MFG_left | 0.54888838 | 0.00045418 | 188.09375 | 821 | 93063.875 |
|  | MFG_right | 0.0509691 | 0.0344719 | 95.6341463 | 387 | 18982.7805 |
|  | OFC_left | 0.00031023 | 0.06921801 | 92.9333333 | 734 | 37196.4167 |
|  | OG_left | 0.9998439 | 0.00158515 | 331.045455 | 960 | 187216.818 |
|  | PMC_left | 0.94790105 | 0.06262299 | 173.590909 | 972 | 73297.0682 |
|  | S1_left | 0.96454817 | 0.22262812 | 122 | 788 | 39404.8571 |
|  | SMA_left | 0.6452346 | 0.01843759 | 351.555556 | 901 | 85933 |
|  | TPJ_right | 0.83032309 | 0.31234647 | 182.375 | 964 | 1224.58333 |
| POPOUT | ACC_right | 0.294054 | 0.409955 | 52.51667 | 311 | 4251.583 |
|  | M1_left | 0.352475 | 0.001038 | 114.2174 | 475 | 33413.28 |
|  | MCC_left | 0.691828 | 0.004027 | 100.25 | 359 | 15481.19 |
|  | MFG_left | 0.965614 | 0.011273 | 134.8125 | 587 | 39769.16 |
|  | MFG_right | 0.065358 | 0.135731 | 49.19512 | 434 | 9291.134 |
|  | OFC_left | 0.010252 | 0.041536 | 115.0167 | 564 | 32087.95 |
|  | OG_left | 0.512003 | 0.051584 | 140.6818 | 465 | 19032.32 |
|  | PMC_left | 0.902396 | 0.003943 | 136.8182 | 746 | 52605.75 |
|  | S1_left | 0.413303 | 0.105759 | 94.28571 | 315 | 13111.04 |
|  | SMA_left | 0.611452 | 0.002316 | 249.1667 | 585 | 91515.72 |
|  | TPJ_right | 0.086352 | 0.073907 | 183.625 | 758 | 88775.33 |

*Highlighted in bright green background are the ROIs in each condition that passed both validity criteria.*

**Supplementary Table 8.** Statistical comparison of within-condition Fast versus Slow DTW alignment across the eleven valid ROIs. Results comparing peak/integral offset against a null distribution, obtained from a within-participant condition-label-shuffle (1,000 permutations; two-sided *p*). *(The table is shown in next page)* *Results exceeding the null at p<0.05 are highlighted in a bright green background for each ROI*.

|  | ROI | Peak<br>(empirical<br>) | Peak<br>(null mean) | Peak<br>(percentile) | Peak<br>(p) | Integral<br>(empirical) | Integral<br>(null mean) | Integral<br>(percentile) | Integral<br>(p) |
| --- | --- | --- | --- | --- | --- | --- | --- | --- | --- |
| S<br>E<br>A<br>R<br>C<br>H | M1_left | 237.97826<br>1 | 55.6517826 | 100 | 0.001 | 68494.3043 | 2364.94554 | 99.1 | 0.001 |
|  | MFG_left | 188.13541<br>7 | 37.1690521 | 100 | 0.001 | 93071.8125 | -1830.4841 | 100 | 0.001 |
|  | OG_left | 331.04545<br>5 | 84.4014091 | 100 | 0.001 | 187216.818 | 6845.19327 | 100 | 0.001 |
|  | PMC_left | 173.77272<br>7 | 60.2399318 | 98.9 | 0.012 | 73328.25 | -1570.275 | 98.9 | 0.012 |
|  | S1_left | 122 | 76.1258571 | 79.9 | 0.202 | 39404.8571 | -515.15032 | 84.6 | 0.155 |
|  | SMA_left | 351.55555<br>6 | 82.6911667 | 100 | 0.001 | 85933 | 292.348056 | 98.9 | 0.012 |
|  | MCC_left | 96.6875 | 50.6675417 | 90.2 | 0.099 | 44358.4583 | -1195.1867 | 96.6 | 0.035 |
|  | ACC_right | 58 | 46.6144 | 67.4 | 0.327 | 3625.43333 | -1277.3019 | 53.4 | 0.466 |
|  | MFG_right | 95.634146<br>3 | 41.9524268 | 96 | 0.041 | 18982.7805 | -22.635915 | 87.1 | 0.129 |
|  | OFC_left | 92.933333<br>3 | 51.3989167 | 87.6 | 0.125 | 37196.4167 | 374.42665 | 94.2 | 0.059 |
|  | TPJ_right | 182.375 | 75.6895833 | 96.5 | 0.036 | 1224.58333 | -1196.113 | 46.4 | 0.536 |
|  | POOLED | 127.70454<br>5 | 17.4437727 | 100 | 0.001 | 60260.4962 | 178.268458 | 100 | 0.001 |
| P<br>O<br>P<br>O<br>U<br>T | M1_left | 114.21739<br>1 | 39.231913 | 99.4 | 0.007 | 33413.2826 | -397.35591 | 99.8 | 0.003 |
|  | MFG_left | 135.22916<br>7 | 29.5826042 | 100 | 0.001 | 39792.9688 | -1323.2988 | 100 | 0.001 |
|  | OG_left | 140.68181<br>8 | 52.0074545 | 98.8 | 0.013 | 19032.3182 | 1693.33791 | 93 | 0.071 |
|  | PMC_left | 136.81818<br>2 | 39.0414773 | 100 | 0.001 | 52734.0682 | -61.754 | 100 | 0.001 |
|  | S1_left | 94.285714<br>3 | 47.9760357 | 91.7 | 0.084 | 13111.0357 | -1312.126 | 84.6 | 0.155 |
|  | SMA_left | 249.16666<br>7 | 59.8576667 | 100 | 0.001 | 91515.7222 | -928.38722 | 100 | 0.001 |
|  | MCC_left | 100.25 | 35.200875 | 98.9 | 0.012 | 15481.1875 | -1970.3986 | 93.3 | 0.068 |
|  | ACC_right | 52.516666<br>7 | 37.6118 | 75.6 | 0.245 | 4251.58333 | -878.8274 | 63.2 | 0.368 |
|  | MFG_right | 49.195122 | 31.3760854 | 78.7 | 0.214 | 9291.13415 | -1547.5275 | 82.8 | 0.173 |
|  | OFC_left | 115.05 | 40.0225667 | 99.4 | 0.007 | 32090.7667 | -561.0557 | 98.9 | 0.012 |
|  | TPJ_right | 183.625 | 67.91375 | 98.7 | 0.014 | 88770.4167 | 1669.01854 | 99.3 | 0.008 |
|  | POOLED | 92.435606<br>1 | 12.4977519 | 100 | 0.001 | 23578.786 | -480.72883 | 100 | 0.001 |

**Supplementary Fig. 6.**
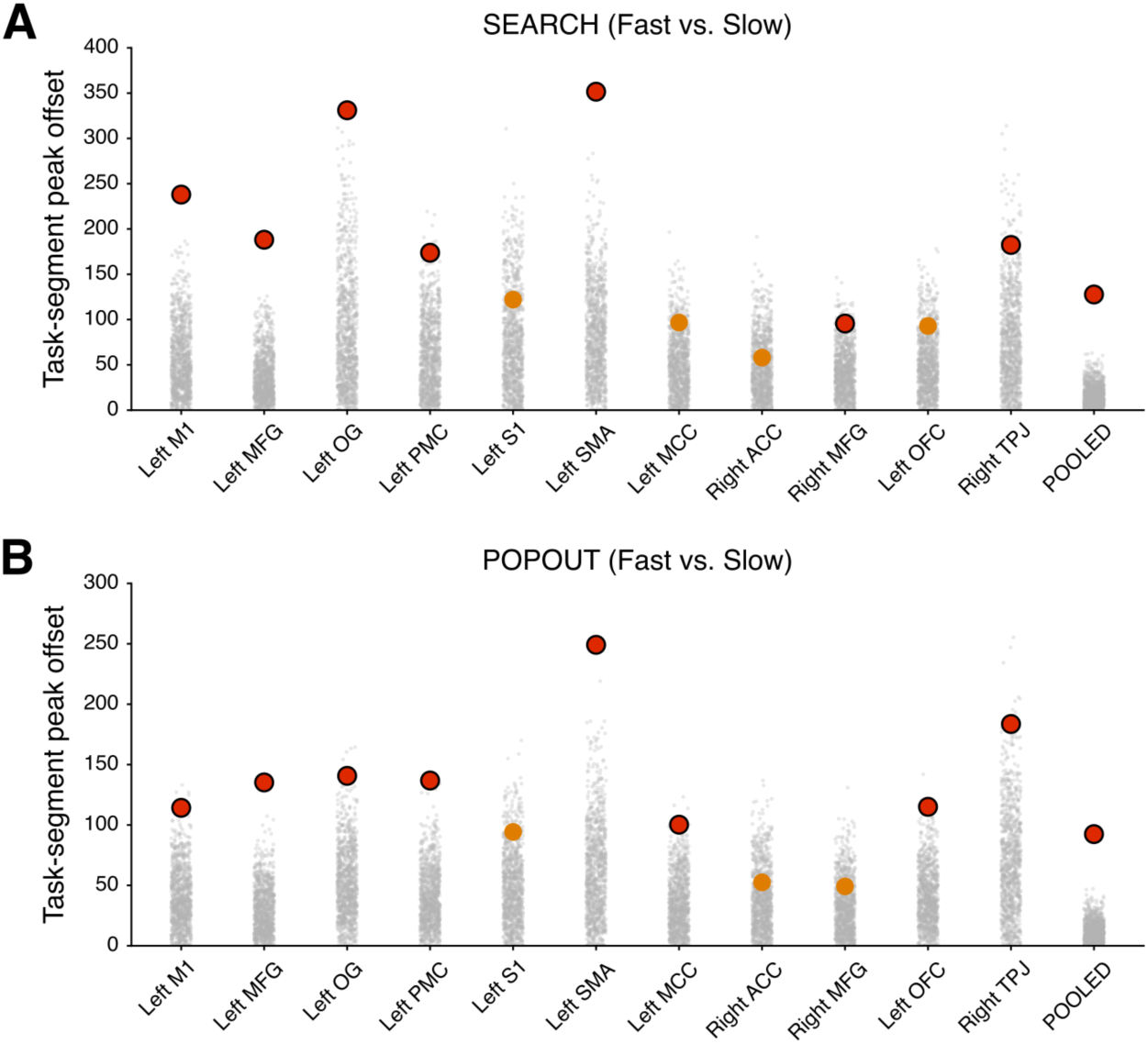
The within-condition analyses comparing Fast versus Slow trials reveal that the temporal scaling is present in each condition. **A**, Task-segment peak warping offset between Fast and Slow Search is shown for each valid ROI comparing the empirical value against a null distribution. **B**, Task-segment peak warping offset between Fast and Slow Pop-out is shown for each valid ROI comparing the empirical value against a null distribution. In each plot, the empirical value for each valid ROI is shown in red (black circled) when exceeding the null (*p*<0.05) or in orange when it did not. Each gray dot represents one permutation defining the null distribution (1,000 permutations).

#### SI – 3.5 Dynamic time warping in control low-frequency bands

The dynamic time warping analysis was repeated in control frequency bands (beta: 14–30 Hz; alpha: 8–13 Hz). Beta-band suppression, in particular, provides an empirical counterpart to a response-bounded null: it is a robustly task-modulated signal (Fig. 1E) bounded by the same stimulus-to-response interval as HFA, in the same regions. If the warping offset were a generic consequence of that interval, beta would exhibit the same scaling. Of the eleven ROIs valid for HFA, none showed a network-wide beta or alpha offset (only Left OG, the primary visual region, survived; Supplementary Table 9), establishing that the effect is specific to HFA rather than a property of any response-bounded signal.

**Supplementary Table 9.**
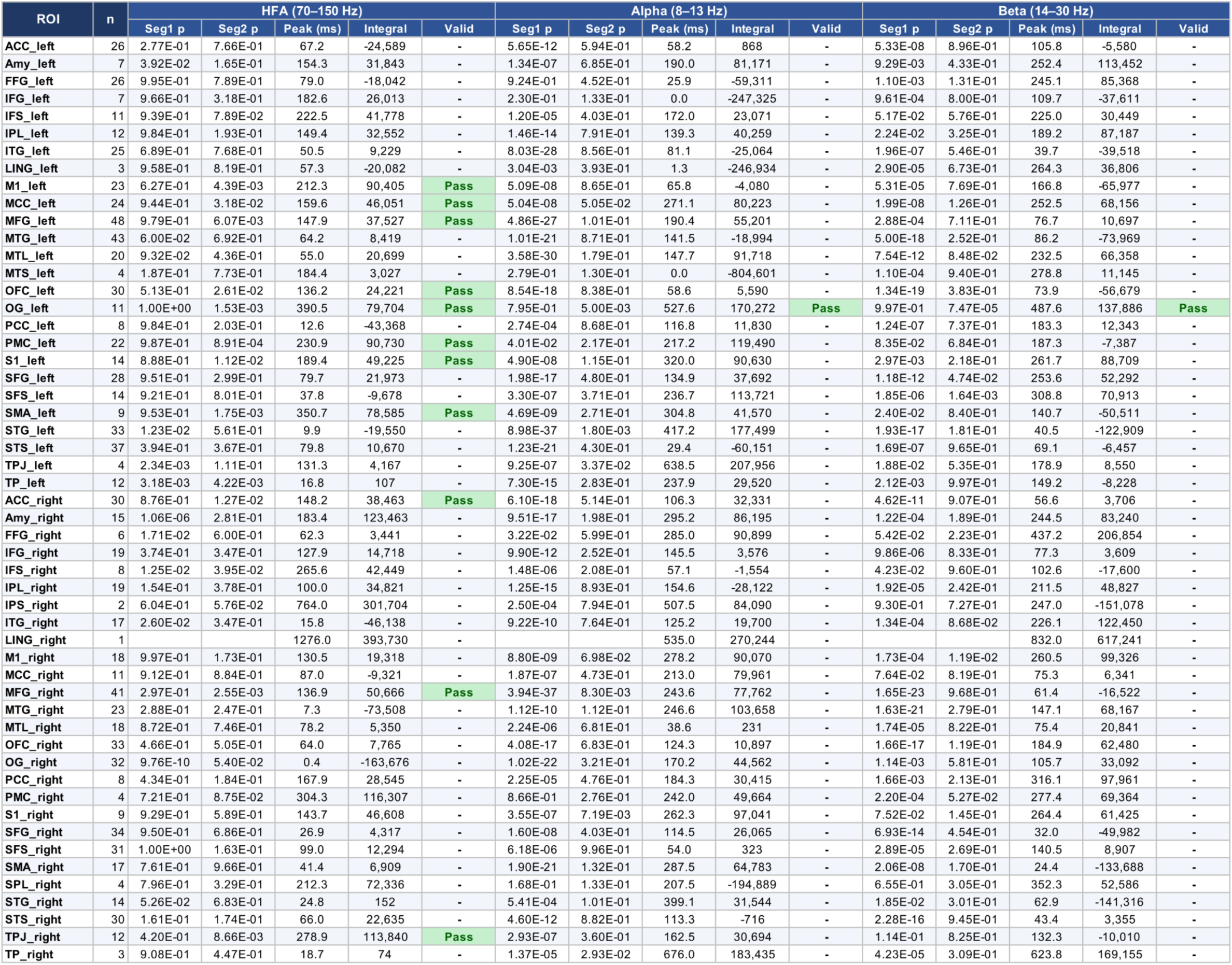
Band-specificity of the temporal stretching effect across ROIs. DTW warping-offset analysis (stimulus-locked) repeated with three frequency bands: HFA, alpha, and beta.

### SI – 4. Phase-amplitude coupling (PAC)

To assess whether alpha–HFA phase-amplitude coupling is specific to visual search or whether it is a fairly inherent property of occipital/parietal cortex, we repeated the phase-amplitude coupling (PAC) analysis within the pre-stimulus interval [-2000, 0 ms] relative to search display onset. The pipeline was identical to the one used for the stimulus-locked interval. The selected interval has the same length as the one used in the main analysis (search window; Fig. 4), and it starts right at fixation onset, just after the ITI, to avoid previous-trial contamination. However, it is worth noting that this window spans the fixation interval (500 ms), sample interval (1,000 ms), and working-memory delay (500 ms). Thus, it cannot be considered a neutral baseline but rather a period of active cognition, encompassing target encoding and short maintenance rather than search itself.

The results in patient IR45 confirmed that HFA amplitude was modulated as a function of alpha phase, in both Search and Pop-out (Supplementary Fig. 7A). The proportion of electrodes with statistically significant PAC (surrogate-based threshold) was highest in posterior regions in both conditions, peaking in OG (Pop-out: 100%; Search: 100%), FFG (83%; 83%), and IPL (57%; 85%), as well as M1 (75%; 75%) and MTL (75%; 25%), and generally declined slightly through PMC (40%; 40%) and S1 (50%; 25%) (Supplementary Fig. 7B). Single-trial PAC estimates confirmed that HFA amplitude was maximal at a preferred alpha phase across individual trials in both Search and Pop-out (Supplementary Fig. 7C). Phase histogram and polar distributions of the preferred alpha phase across significant channels clustered at the same angular location in both conditions. The preferred phase angle (∼160°–180°) was the same as in the main analysis in the search display post-stimulus interval [0, +2000 ms]. These results indicate that alpha– HFA coupling is not specific to the visual-search period: the same coupling, at the same preferred phase, is already present during target encoding and maintenance. This identifies alpha-phase gating of HFA as a general property of the posterior cortex engaged across task phases, rather than a mechanism evoked specifically by visual search.

**Supplementary Fig. 7.**
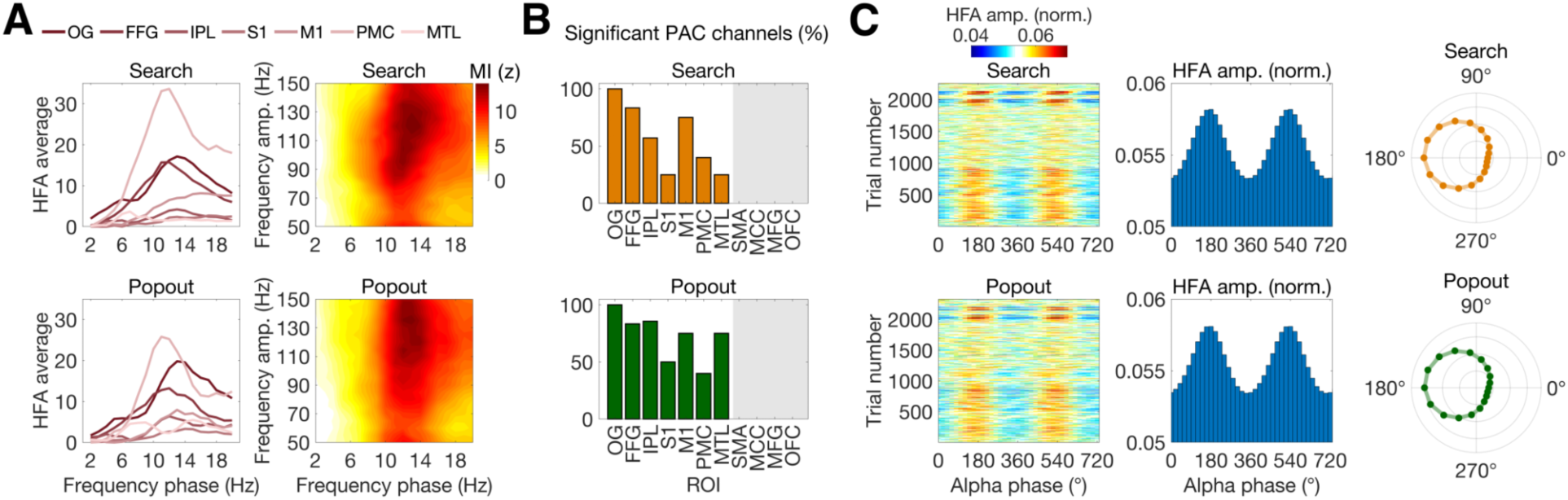
Alpha–HFA phase-amplitude coupling is present also before stimulus presentation. Single-participant data (patient *IR45*; the only participant with Left OG coverage) shown as in Fig. 4. **A**, Left: mean HFA amplitude as a function of low-frequency phase for each ROI (color-coded lines; see legend), for Search (top) and Pop-out (bottom). Posterior regions show prominent sinusoidal modulation of HFA amplitude as a function of alpha phase. Right: grand-average frequency-frequency modulation index (MI, z-scored against surrogate distributions) comodulograms, showing coupling between phase frequencies (x-axis, 2–20 Hz) and amplitude frequencies (y-axis, 50–150 Hz). A prominent alpha–HFA coupling peak (∼8–12 Hz phase, 70–150 Hz amplitude) is present in both conditions. **B**, Percentage of electrodes with statistically significant PAC (*z*>2.33, *p*<0.01, surrogate-based threshold) per ROI, for Search (orange) and Pop-out (green). **C**, Single-trial alpha–HFA PAC in the ROIs shown in B. Left: HFA amplitude as a function of alpha phase for individual trials (y-axis: trial number). Center: trial-and-channel-averaged phase histogram (bar plots), showing mean HFA amplitude as a function of alpha-band phase (0°–720°, two full cycles shown for clarity). Right: preferred alpha phase for each significant channel (polar plots). HFA amplitude is consistently maximal at a preferred alpha phase in both Search (orange) and Pop-out (green), with phase preferences clustering tightly across channels and trials.

## References

1. Awh, E., Belopolsky, A. V., & Theeuwes, J. (2012). Top-down versus bottom-up attentional control: A failed theoretical dichotomy. Trends in Cognitive Sciences, 16(8), 437–443. 10.1016/j.tics.2012.06.010

2. Buschman, T. J., & Miller, E. K. (2007). Top-Down Versus Bottom-Up Control of Attention in the Prefrontal and Posterior Parietal Cortices. Science, 315(5820), 1860–1862. 10.1126/science.1138071

3. Buschman, T. J., & Miller, E. K. (2009). Serial, Covert Shifts of Attention during Visual Search Are Reflected by the Frontal Eye Fields and Correlated with Population Oscillations. Neuron, 63(3), 386–396. 10.1016/j.neuron.2009.06.020

4. Buzsáki, G., & Draguhn, A. (2004). Neuronal Oscillations in Cortical Networks. Science, 304(5679), 1926– 1929. 10.1126/science.1099745

5. Cohen, M. X. (2014). Analyzing Neural Time Series Data: Theory and Practice. MIT Press.

6. Cohen, M. X. (2017). Where Does EEG Come From and What Does It Mean? Trends in Neurosciences, 40(4), 208–218. 10.1016/j.tins.2017.02.004

7. Corbetta, M., Patel, G., & Shulman, G. L. (2008). The Reorienting System of the Human Brain: From Environment to Theory of Mind. Neuron, 58(3), 306–324. 10.1016/j.neuron.2008.04.017

8. Corbetta, M., & Shulman, G. L. (2002). Control of goal-directed and stimulus-driven attention in the brain. Nature Reviews Neuroscience, 3(3), 201–215. 10.1038/nrn755

9. Daume, J., Kamiński, J., Schjetnan, A. G. P., Salimpour, Y., Khan, U., Kyzar, M., Reed, C. M., Anderson, W. S., Valiante, T. A., Mamelak, A. N., & Rutishauser, U. (2024). Control of working memory by phase–amplitude coupling of human hippocampal neurons. Nature, 1–9. 10.1038/s41586-024-07309-z

10. De Houwer, J. (2019). Moving beyond System 1 and System 2: Conditioning, implicit evaluation, and habitual responding might be mediated by relational knowledge. Experimental Psychology, 66(4), 257–265. 10.1027/1618-3169/a000450

11. Doyon, J., Bellec, P., Amsel, R., Penhune, V., Monchi, O., Carrier, J., Lehéricy, S., & Benali, H. (2009). Contributions of the basal ganglia and functionally related brain structures to motor learning. *Behavioural Brain Research*, Special Issue on the Role of the Basal Ganglia in Learning and Memory, 199(1), 61–75. 10.1016/j.bbr.2008.11.012

12. Duncan, J., & Humphreys, G. (1992). Beyond the search surface: Visual search and attentional engagement. Journal of Experimental Psychology. Human Perception and Performance, 18(2), 578–588; discussion 589-593. 10.1037//0096-1523.18.2.578

13. Duncan, J., & Humphreys, G. W. (1989). Visual search and stimulus similarity. Psychological Review, 96(3), 433–458. 10.1037/0033-295X.96.3.433

14. Eckstein, M. P. (2011). Visual search: A retrospective. Journal of Vision, 11(5), 14. 10.1167/11.5.14

15. Eckstein, M. P. (2017). Probabilistic Computations for Attention, Eye Movements, and Search. Annual Review of Vision Science, 3, 319–342. 10.1146/annurev-vision-102016-061220

16. Engel, A. K., & Fries, P. (2010). Beta-band oscillations—Signalling the status quo? Current Opinion in Neurobiology, 20(2), 156–165. 10.1016/j.conb.2010.02.015

17. Evans, J. St. B. T., & Stanovich, K. E. (2013). Dual-Process Theories of Higher Cognition: Advancing the Debate. Perspectives on Psychological Science, 8(3), 223–241. 10.1177/1745691612460685

18. Fodor, J. A. (1983). The Modularity of Mind. The MIT Press. 10.7551/mitpress/4737.001.0001

19. Fries, P. (2005). A mechanism for cognitive dynamics: Neuronal communication through neuronal coherence. Trends in Cognitive Sciences, 9(10), 474–480. 10.1016/j.tics.2005.08.011

20. Fries, P. (2015). Rhythms for Cognition: Communication through Coherence. Neuron, 88(1), 220–235. 10.1016/j.neuron.2015.09.034

21. Geweke, J. F. (1982). Measurement of linear dependence and feedback between multiple time series. Journal of the American Statistical Association, 77(378), 304–313. 10.2307/2287238

22. Gold, J. I., & Shadlen, M. N. (2007). The Neural Basis of Decision Making. Annual Review of Neuroscience, 30(1), 535–574. 10.1146/annurev.neuro.29.051605.113038

23. Jensen, O., & Mazaheri, A. (2010). Shaping Functional Architecture by Oscillatory Alpha Activity: Gating by Inhibition. Frontiers in Human Neuroscience, 4, 186. 10.3389/fnhum.2010.00186

24. Kahneman, D. (2011). Thinking, fast and slow (p. 499). Farrar, Straus and Giroux.

25. Katsuki, F., & Constantinidis, C. (2014). Bottom-Up and Top-Down Attention: Different Processes and Overlapping Neural Systems. The Neuroscientist, 20(5), 509–521. 10.1177/1073858413514136

26. Klimesch, W. (2012). Alpha-band oscillations, attention, and controlled access to stored information. Trends in Cognitive Sciences, 16(12), 606–617. 10.1016/j.tics.2012.10.007

27. Lei, T., Scheid, M. R., Flint, R. D., Glaser, J. I., & Slutzky, M. W. (2026). Active dissociation of intracortical spiking and high gamma activity. Nature, 1–7. 10.1038/s41586-026-10331-y

28. Leonard, M. K., Gwilliams, L., Sellers, K. K., Chung, J. E., Xu, D., Mischler, G., Mesgarani, N., Welkenhuysen, M., Dutta, B., & Chang, E. F. (2023). Large-scale single-neuron speech sound encoding across the depth of human cortex. Nature, 1–10. 10.1038/s41586-023-06839-2

29. Leszczyński, M., Barczak, A., Kajikawa, Y., Ulbert, I., Falchier, A. Y., Tal, I., Haegens, S., Melloni, L., Knight, R. T., & Schroeder, C. E. (2020). Dissociation of broadband high-frequency activity and neuronal firing in the neocortex. Science Advances, 6(33), eabb0977. 10.1126/sciadv.abb0977

30. Li, L., Gratton, C., Yao, D., & Knight, R. T. (2010). Role of frontal and parietal cortices in the control of bottom-up and top-down attention in humans. Brain Research, 1344, 173–184. 10.1016/j.brainres.2010.05.016

31. Logothetis, N. K. (2008). What we can do and what we cannot do with fMRI. Nature, 453(7197), 869–878. 10.1038/nature06976

32. Orsten-Hooge, K. D., Portillo, M. C., & Pomerantz, J. R. (2015). False pop out. Journal of Experimental Psychology: Human Perception and Performance, 41(6), 1623–1633. 10.1037/xhp0000077

33. Ossandón, T., Vidal, J. R., Ciumas, C., Jerbi, K., Hamamé, C. M., Dalal, S. S., Bertrand, O., Minotti, L., Kahane, P., & Lachaux, J.-P. (2012). Efficient “Pop-Out” Visual Search Elicits Sustained Broadband Gamma Activity in the Dorsal Attention Network. Journal of Neuroscience, 32(10), 3414–3421. 10.1523/JNEUROSCI.6048-11.2012

34. Packard, M. G., & Goodman, J. (2013). Factors that influence the relative use of multiple memory systems. Hippocampus, 23(11), 1044–1052. 10.1002/hipo.22178

35. Pagnotta, M. F., Dhamala, M., & Plomp, G. (2018a). Assessing the performance of Granger–Geweke causality: Benchmark dataset and simulation framework. Data in Brief, 21, 833–851. 10.1016/j.dib.2018.10.034

36. Pagnotta, M. F., Dhamala, M., & Plomp, G. (2018b). Benchmarking nonparametric Granger causality: Robustness against downsampling and influence of spectral decomposition parameters. NeuroImage, 183, 478–494. 10.1016/j.neuroimage.2018.07.046

37. Pagnotta, M. F., Pascucci, D., & Plomp, G. (2020). Nested oscillations and brain connectivity during sequential stages of feature-based attention. NeuroImage, 223, 117354. 10.1016/j.neuroimage.2020.117354

38. Pagnotta, M. F., Pascucci, D., & Plomp, G. (2022). Selective attention involves a feature-specific sequential release from inhibitory gating. NeuroImage, 246, 118782. 10.1016/j.neuroimage.2021.118782

39. Parvizi, J., & Kastner, S. (2018). Promises and limitations of human intracranial electroencephalography. Nature Neuroscience, 21(4), 474–483. 10.1038/s41593-018-0108-2

40. Pracar, A. L., Pagnotta, M. F., Quiroga-Martinez, D. R., Ghuman, R. S., Du, C., Dastjerdi, M., Lin, J. J., Willie, J. T., Brunner, P., Dronkers, N. F., & Knight, R. T. (2026). Sensorimotor dynamics differentiate singing and speaking (p. 2025.12.19.695607). bioRxiv. 10.64898/2025.12.19.695607

41. Ray, S., & Maunsell, J. H. R. (2011). Different Origins of Gamma Rhythm and High-Gamma Activity in Macaque Visual Cortex. PLOS Biology, 9(4), e1000610. 10.1371/journal.pbio.1000610

42. Sakoe, H., & Chiba, S. (1978). Dynamic programming algorithm optimization for spoken word recognition. IEEE Transactions on Acoustics, Speech, and Signal Processing, 26(1), 43–49. 10.1109/TASSP.1978.1163055

43. Slama, S. J. K., Jimenez, R., Saha, S., King-Stephens, D., Laxer, K. D., Weber, P. B., Endestad, T., Ivanovic, J., Larsson, P. G., Solbakk, A.-K., Lin, J. J., & Knight, R. T. (2021). Intracranial Recordings Demonstrate Both Cortical and Medial Temporal Lobe Engagement in Visual Search in Humans. Journal of Cognitive Neuroscience, 33(9), 1833–1861. 10.1162/jocn_a_01739

44. Squire, L. R., & Dede, A. J. O. (2015). Conscious and Unconscious Memory Systems. Cold Spring Harbor Perspectives in Biology, 7(3), a021667. 10.1101/cshperspect.a021667

45. Tavenard, R., Faouzi, J., Vandewiele, G., Divo, F., Androz, G., Holtz, C., Payne, M., Yurchak, R., Rußwurm, M., Kolar, K., & Woods, E. (2020). Tslearn, A Machine Learning Toolkit for Time Series Data. Journal of Machine Learning Research, 21(118), 1–6.

46. Tort, A. B. L., Komorowski, R., Eichenbaum, H., & Kopell, N. (2010). Measuring Phase-Amplitude Coupling Between Neuronal Oscillations of Different Frequencies. Journal of Neurophysiology, 104(2), 1195–1210. 10.1152/jn.00106.2010

47. Treisman, A. M., & Gelade, G. (1980). A feature-integration theory of attention. Cognitive Psychology, 12(1), 97–136. 10.1016/0010-0285(80)90005-5

48. Treisman, A., & Sato, S. (1990). Conjunction search revisited. Journal of Experimental Psychology: Human Perception and Performance, 16(3), 459–478. 10.1037/0096-1523.16.3.459

49. Varela, F., Lachaux, J.-P., Rodriguez, E., & Martinerie, J. (2001). The brainweb: Phase synchronization and large-scale integration. Nature Reviews Neuroscience, 2(4), 229–239. 10.1038/35067550

50. Vintsyuk, T. K. (1968). Speech discrimination by dynamic programming. Cybernetics, 4(1), 52–57. 10.1007/BF01074755

51. Wolfe, J. M. (2014). Approaches to visual search: Feature integration theory and guided search. In The Oxford handbook of attention (pp. 11–55). Oxford University Press. 10.1093/oxfordhb/9780199675111.013.002

52. Wolfe, J. M. (2021). Guided Search 6.0: An updated model of visual search. Psychonomic Bulletin & Review, 28(4), 1060–1092. 10.3758/s13423-020-01859-9

53. Wolfe, J. M., Friedman-Hill, S. R., Stewart, M. I., & O’Connell, K. M. (1992). The role of categorization in visual search for orientation. Journal of Experimental Psychology: Human Perception and Performance, 18(1), 34–49. 10.1037/0096-1523.18.1.34

54. Worden, M. S., Foxe, J. J., Wang, N., & Simpson, G. V. (2000). Anticipatory Biasing of Visuospatial Attention Indexed by Retinotopically Specific α-Bank Electroencephalography Increases over Occipital Cortex. The Journal of Neuroscience, 20(6), RC63–RC63. 10.1523/JNEUROSCI.20-06-j0002.2000

